# Differences in substrate engagement and Retinoblastoma protein (RB) binding of human KDM5A and KDM5B

**DOI:** 10.64898/2026.04.30.721888

**Authors:** Till L. Ruengeler, Egor A. Pavlenko, Fabian Basler, Juliane Renn, Farnusch Kaschani, Marie-Anne Derichs, Linda C. Zirden, Batool Shannan, Alexandra Hommel, Markus Kaiser, Alexander Roesch, Simon Poepsel

## Abstract

Trimethylation of lysine 4 of histone H3 (H3K4me3) is a post-translational modification (PTM) enriched at promoters of actively transcribed genes. H3K4me3 is removed by the human histone demethylases of the KDM5 family. KDM5 demethylases act as transcriptional repressors through their catalytic activity in addition to more complex roles that depend on their interactions with other chromatin regulators and may be independent of demethylase activity. To better understand the mechanistic differences of the closely related paralogs KDM5A and KDM5B as well as their interactions with Retinoblastoma protein (RB), we systematically analyzed and compared their demethylase activities, nucleosome engagement, and RB binding. We used recombinant nucleosome binding and demethylase activity assays, as well as an integrative structural biology approach using negative-stain electron microscopy (EM), AlphaFold predictions, and cross-linking mass spectrometry for a comprehensive *in vitro* analysis of these critical and largely non-redundant enzymes.

KDM5A and KDM5B showed differences in enzyme kinetics using peptide substrates, as well as in nucleosome binding. Furthermore, KDM5A interacts with RB, mainly mediated by its canonical LxCxE RB binding motif. KDM5B, on the other hand, lacks an LxCxE binding motif and does not stably bind to RB under the conditions tested here. RB directly interacts with nucleosomes, and its nucleosome binding does not measurably affect KDM5A demethylase activity or nucleosome interactions. Our findings provide a biochemical framework for the differences between KDM5A and KDM5B regarding RB interactions and nucleosome engagement.

## Introduction

Histone proteins are subject to dynamic post-translational modifications (histone PTMs) that are tightly linked to functional states of chromatin.^[1,2]^ Trimethylation of lysine 4 of histone H3 (H3K4me3) is enriched at the promoters of actively transcribed genes^[3,4]^ and can be reversed by KDM5 histone lysine demethylases^[5,6]^ that belong to the Jumonji C domain family of Fe(II)-and α-ketoglutarate dependent dioxygenases.^[5,7–9]^ There are four human KDM5 demethylases: KDM5A (Jarid1A, RBP2), KDM5B (Jarid1B, PLU1), KDM5C (Jarid1C, SMCX), and KDM5D (Jarid1D, SMCY).^[8]^ They are characterized by a conserved multi-domain architecture, including the catalytic split Jumonji domain, composed of the N- and C-terminal Jumonji domains (JmjN and JmjC, respectively), interspersed with a Plant Homeodomain zinc finger (PHD1) and an AT-rich interaction domain (ARID) (Fig. 1A).^[10–12]^ A C5HC2 zinc finger domain was found to be essential for demethylase activity.^[13]^. C-terminal of the catalytic core, KDM5A and B have two additional PHD domains (PHD2/3), separated from the active core by a conserved spectrin-like, coiled-coil region (Suppl. Fig. 1A,B).^[8,14]^ With the exception of PHD1 and PHD3 in KDM5A/B, no interaction partners have been unambiguously identified and mapped to KDM5 domains, yet.^[8]^ PHD1 binding to unmethylated H3K4 promotes demethylation, representing a positive feedback mechanism.^[14–16]^ PHD3 was shown to preferentially bind to H3K4me3 and may be involved in the targeting of the enzyme. However, the contribution of these domains to specificity or recruitment remains poorly understood in the cellular context.^[16,17]^ The ARID domains of KDM5 demethylases bind to GC-rich DNA, and KDM5 proteins were reported to be enriched at CpG-islands in the genome.^[18–20]^ KDM5A and KDM5B are ubiquitously expressed during development, but KDM5B is poorly expressed in adult tissue except for male testes, while KDM5A is continuously expressed in most tissues, especially in bone tissue.^[21–23]^ KDM5A/B have the highest sequence similarity within the KDM5 family, have overlapping yet distinct biological roles, and are frequently dysregulated in disease.^[23]^ Both are essential for normal tissue development and play roles in DNA damage repair and cell proliferation.^[23]^ Knock-out of KDM5B in mice leads to prenatal mortality,^[24]^ while mutations of KDM5A in humans can lead to neurodevelopmental disorders like autism spectrum disorder.^[25]^ KDM5A and B can partially rescue the loss of the other homolog in some experimental systems ^[20]^, but the two proteins are not functionally redundant. For example, during neural development, KDM5A suppresses neuronal cell fate by repressing Gfap promoters^[26]^, whereas KDM5B promotes neuronal differentiation by repressing neural stem cell markers.^[27]^ In musculoskeletal differentiation, their roles are also distinct. KDM5B maintains an undifferentiated state by suppressing differentiation regulators^[28]^, while KDM5A interacts with the retinoblastoma protein (RB) to promote myoblast differentiation.^[29]^ It has been hypothesized that these functional differences arise from different target genes and protein interactions, but the underlying mechanisms remain unclear.

**Figure 1:**
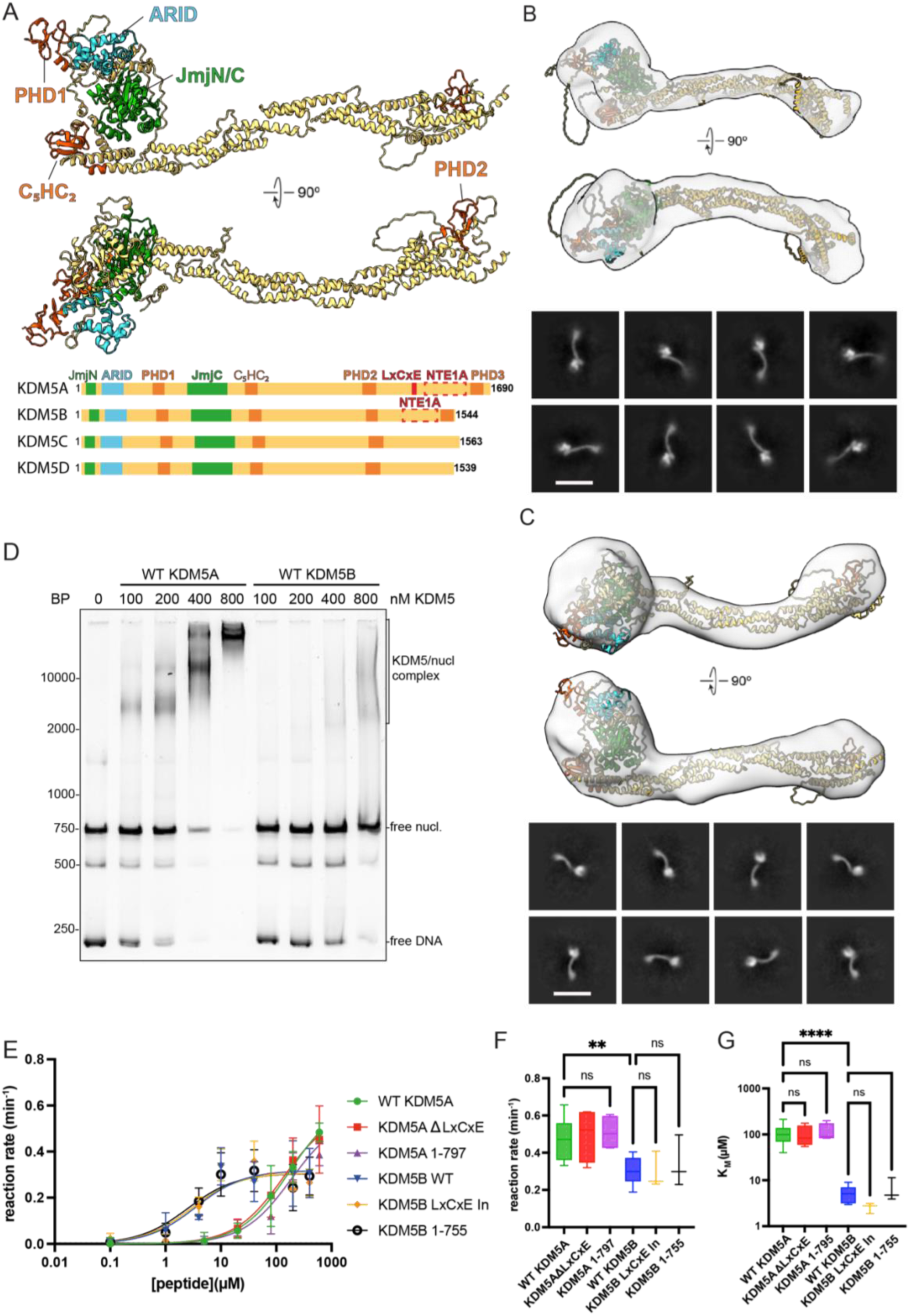
Structure and activity of human KDM5A and. **B.** (A) Top: AlphaFold3 prediction of KDM5B. Regions predicted with low confidence and high disorder content are not shown for clarity (aa234-309; aa1284-1344; aa1370-1562; for the prediction metrics see Suppl. Fig. 3). Bottom: Domain architecture of KDM5 proteins according to Uniprot database annotation. Green, catalytic JmjN/C domain; blue, ARID domain; orange, PHD1-3 and C5HC2 zinc finger domains; red, reported RB interaction sites. (B,C) Negative-stain EM 3D reconstructions (top) and representative 2D class averages (bottom) of KDM5A (B) and KDM5B (C). AlphaFold3 predictions were fitted into the densities by rigid-body docking. The AlphaFold3 models were separated after aa797 for KDM5A and after aa755 for KDM5B to accommodate the orientation of the C-terminal coiled-coil region. Scale bars = 24 nm. (D) Electrophoretic mobility shift assay (EMSA) of KDM5A and KDM5B with 100 nM H3K4me3MLA nucleosomes (nucl). DNA stained with SybrGold. BP: base pairs (marker). Image representative of five independent experiments (E-G). FDH-based demethylase activity assays with H3_1-18_K4me3 peptides. Michaelis–Menten plot (F) and respective calculated kinetic rate in min^-1^ (G) and K_M_ in µM (H) of WT KDM5A (green), KDM5A **Δ**LxCxE (red), KDM5A active core (KDM5A 1-797), (purple), WT KDM5B (blue), KDM5B LxCxE insertion, and KDM5B active core (KDM5B1-755). Means of at least three independent experiments and the standard error of the mean (SEM) are shown. Significance was determined using one-way ANOVA, with P < 0.05 considered significant. P values below 0.01 are highlighted with two asterisks and below 0.0001 with four.

The first reported interactor of KDM5 demethylases is RB ^[30]^, a co-regulator of E2F transcription factors that performs many and complex functions in cell cycle progression, senescence, differentiation, and DNA damage repair.^[31,32]^ A central mechanism of transcription regulation by RB is based on its interaction with transcription factors, which is dependent on RB phosphorylation.^[33,34]^ In addition to its role as a cell cycle regulator performed primarily by suppressing E2F target genes, RB has also been reported to interact with chromatin regulators such as enhancer of zeste homolog 2 (EZH2) and the nucleosome-remodeling deacetylase (NuRD) complex during replication and to facilitate DNA repair.^[35,36]^ Several studies have investigated the functional interplay between KDM5 demethylases and RB. For example, RB was reported to recruit KDM5 proteins to E2F target genes to repress their expression, possibly supported by demethylation of H3K4me3, ultimately resulting in cell cycle arrest and senescence.^[37–39]^ During differentiation, RB coordinates with KDM5A and E2F4 to repress E2F4/RB target genes while activating transcription of differentiation genes to maintain a differentiated cell state.^[39,40]^ Several studies report that RB deficient cancer cells are sensitive to the experimental modulation of KDM5 function. For example, KDM5A knock-out in RB deficient mice suppresses tumor formation ^[41]^, and in RB deficient osteosarcoma or retinoblastoma, overexpression of KDM5A reduces tumor growth by regulating mitochondrial function.^[29]^ These observations imply that the effects of KDM5A RB interactions depend on the cellular context. A similar functional relationship has also been observed for KDM5B and RB in melanoma cells, where KDM5B stabilizes hypophosphorylation of serine 795 of RB, which is associated with cell cycle control and thus is involved in suppression of tumor growth and therapy resistance.^[42,43]^ To better determine whether KDM5 demethylases are relevant therapeutic targets in RB deficient cancers, the molecular basis of their interaction with RB, and especially the differences between KDM5A and KDM5B in this context, must be studied in detail.^[23]^

KDM5A has been reported to bind RB via two interfaces: an LxCxE motif, also used by viral oncoproteins to target RB, including papillomaviral proteins^[44,45]^, and an additional non-T/E1A region (NTE1A), located C-terminal to the LxCxE motif.^[46,47]^ RB binding to the LxCxE motif is well characterized for other binding partners. The LxCxE motif binds to a solvent-exposed region of RB pocket B (RB pB), which, together with RB pocket A (RB pA), forms the large pocket domain (RB LP), the main interaction site of RB.^[44]^ KDM5B was also identified as interacting with RB in co-IP experiments, but lacks an LxCxE motif, containing only an NTE1A region.^[48]^

Although KDM5A and B show the same biochemical activity and share a highly similar domain architecture, yet they are not functionally redundant. Dissecting molecular mechanisms of their shared yet divergent properties requires rigorous analysis grounded in biochemical and structural data. Here, we investigate key properties of KDM5A and KDM5B *in vitro*, with a focus on their topology, nucleosome binding, histone demethylase activity, and their interactions with RB. While their overall structures are comparable at low resolution, both enzymes show divergent catalytic properties towards histone peptide substrates and differ in nucleosome binding in isolation. KDM5A, but not KDM5B, can stably interact with recombinant human RB *in vitro*, and this interaction is dependent on RB pocket interactions of the LxCxE motif that is found exclusively in KDM5A. KDM5A can bind to and demethylate histone peptides and nucleosomes independently of RB. Our work provides a framework for understanding and further investigating the mechanistic differences between KDM5A and B and the molecular implications of RB binding.

## Results

### Structures and catalytic activity of KDM5A and KDM5B

All four human KDM5 demethylases share a similar domain arrangement (Fig. 1A). At the amino acid (aa) sequence level, KDM5A and B are most closely related in their JmjC, JmjN, and PHD1 domains (with 87 %, 79 %, and 84 % sequence identity respectively), while the largest divergence in sequence is found in the regions between the ARID and PHD1 domains, and between PHD2 and PHD3 (Suppl. Fig. 1A,B). The region between the ARID and PHD1 domains constitutes an intrinsically disordered region (IDR) that was reported to mediate nucleosome interactions in the case of KDM5A (Suppl. Fig. 1C).^[49]^ The region between PHD2 and 3 is also disordered and encompasses potential protein-protein interaction sites, including the LxCxE motif and the NTE1A region, which reportedly mediate RB binding.^[45–47]^ Thus, the aa sequences of KDM5A and B differ most strongly in disordered regions that are likely of functional importance.

Since high-resolution structural data on KDM5 demethylases remain scarce and limited to the catalytically active core, we performed single-particle electron microscopy (EM) studies to investigate the topologies of full-length human KDM5A and KDM5B. Attempts to obtain cryogenic electron microscopy (cryo-EM) reconstructions were unsuccessful, likely due to the low thermal stability of the full-length proteins (data not shown) and high denaturation propensity during the vitrification process. We therefore performed negative-stain EM analyses of recombinant human, full-length KDM5A and KDM5B (Suppl. Fig. 2A,B, 3,4). Within the resolution limits of negative-stain EM, both proteins displayed highly similar overall shapes and architectures (Fig. 1B,C). AlphaFold models of full-length KDM5A and KDM5B (Suppl. Fig. 5) were rigid-body fitted into the corresponding 3D reconstruction with only minor adjustments of the orientation of the spectrin-like, coiled-coil region (Fig. 1B,C). In both cases, the N-terminal catalytic module forms a compact core, while the C-terminal part, comprising the spectrin-like, coiled-coil region and PHD2/3, extends as an elongated structure from the core (Fig. 1 B,C). This arrangement is in agreement with previous reports on negative-stain reconstructions of KDM5B.^[50]^ Notably, the varying angles at which the elongated, C-terminal parts of KDM5A and B extend from the catalytic core (Fig. 1B,C) indicate a degree of relative flexibility in this region. The C-terminal PHD3 domain preferentially binds to H3K4me3^[16,17]^, is separated from the remainder of the proteins by an unstructured linker, and was not predicted with confidence to interact with other parts of KDM5A/B (Suppl. Fig. 5), leaving open the question of whether this domain participates in stable intramolecular interactions or whether and how it may be involved in substrate recognition.

To analyze whether the global structural similarity between KDM5A and B is mirrored at the functional level, we performed nucleosome-binding and histone demethylation assays for both enzymes under identical conditions. Substrate engagement was monitored using recombinant nucleosomes bearing pseudo-trimethylated H3K4 (H3K4me3MLA)^[51]^ in an electrophoretic mobility shift assay (EMSA). For KDM5A, we observed a decrease in the free nucleosome bands at lower enzyme concentrations, and more defined nucleosome-KDM5 complex bands as compared to KDM5B. This observation indicates more stable and higher-affinity nucleosome binding of KDM5A versus KDM5B (Fig. 1D). To assess enzyme kinetics, we performed formaldehyde dehydrogenase (FDH) coupled activity assays^[14]^ with H3K4me3-bearing histone H3 aa1-18 peptides (H3_1-18_K4me3). Our kinetic analysis showed a higher reaction rate (k_cat_) for KDM5A (0.47±0.11 min^-1^) than for KDM5B (0.3±0.08 min^-1^), whereas KDM5B had a higher substrate affinity than KDM5A (K_M_ of 5.3±2.2 µM and 106.9±49.7 µM, respectively) (Fig. 1E-G, Table 1). Similar differences were found using KDM5 constructs containing only the catalytic core (KDM5A 1-797 and KDM5B 1-755) (Fig. 1E-G, Table 1) and were independently confirmed by a luminescence-based demethylase assay (Suppl. Fig. 2C,D). Thus, while their overall domain architecture seems largely similar, KDM5A and KDM5B display distinct kinetic features that may underlie functional differences and non-redundant functions in cells.

**Table: 1.**
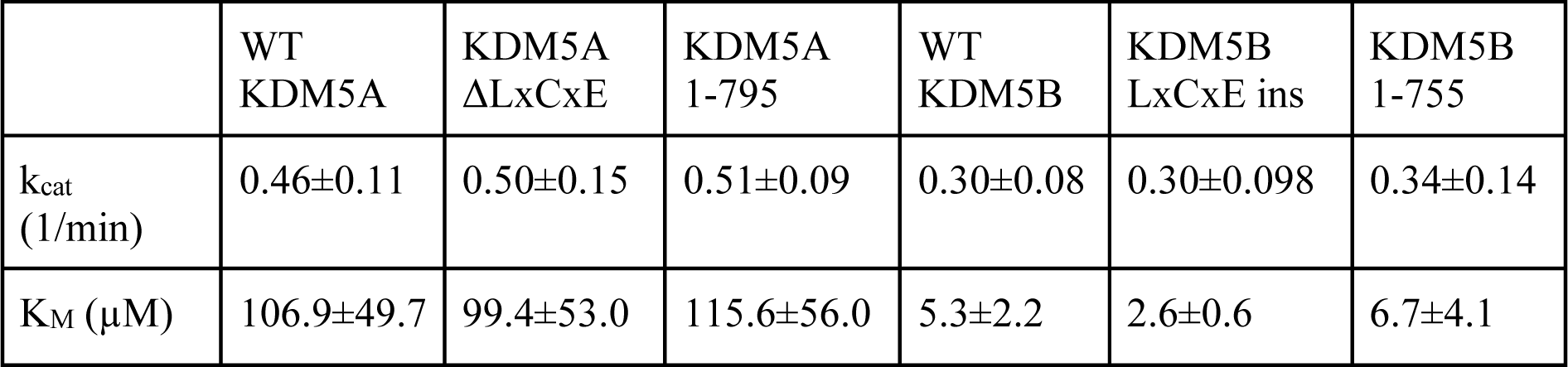
Table showing the k_cat_ and K_M_ of the KDM5 construct obtained from FDH coupled assays with histone H3_1-18_K4me3 peptides as substrate. Means of at least three experiments and the Standard Error of the mean are shown.

| | WT<br>KDM5A | KDM5A<br>ΔLxCxE | KDM5A<br>1-795 | WT<br>KDM5B | KDM5B<br>LxCxE ins | KDM5B<br>1-755 |
| --- | --- | --- | --- | --- | --- | --- |
| $k_{cat}$<br>(1/min) | 0.46±0.11 | 0.50±0.15 | 0.51±0.09 | 0.30±0.08 | 0.30±0.098 | 0.34±0.14 |
| $K_M$ (μM) | 106.9±49.7 | 99.4±53.0 | 115.6±56.0 | 5.3±2.2 | 2.6±0.6 | 6.7±4.1 |

### KDM5 RB interaction

While both KDM5A and B have been described as RB-interacting proteins, only KDM5A contains a canonical RB-binding LxCxE motif that is located in a disordered region N-terminal of the PHD3 domain (Fig. 1A and Suppl. Fig. 1C). KDM5B association with RB has been reported in cells overexpressing different KDM5B truncation variants, narrowing down the RB interactions to the NTE1A region.^[38,42,48]^ To characterize how KDM5A and KDM5B interact with RB, we performed a series of *in vitro* binding experiments using recombinant full-length RB (Suppl. Fig. 6). Given that KDM5A and B show unusual separation behavior in size exclusion chromatography (Suppl. Fig. 2A), we used chemical cross-linking and denaturing SDS-PAGE to assess their interaction with RB. Under these conditions, KDM5A formed a clear cross-linked complex with RB, while KDM5B exhibited no complex band (Fig 2A). These results were corroborated by native PAGE and biolayer interferometry (BLI) (Fig. 2B,C), which together suggest that KDM5B, if at all, interacts only weakly with RB under the investigated conditions. The canonical LxCxE motif present in KDM5A but not KDM5B may therefore be a key determinant of the KDM5A RB interactions.

**Figure 2:**
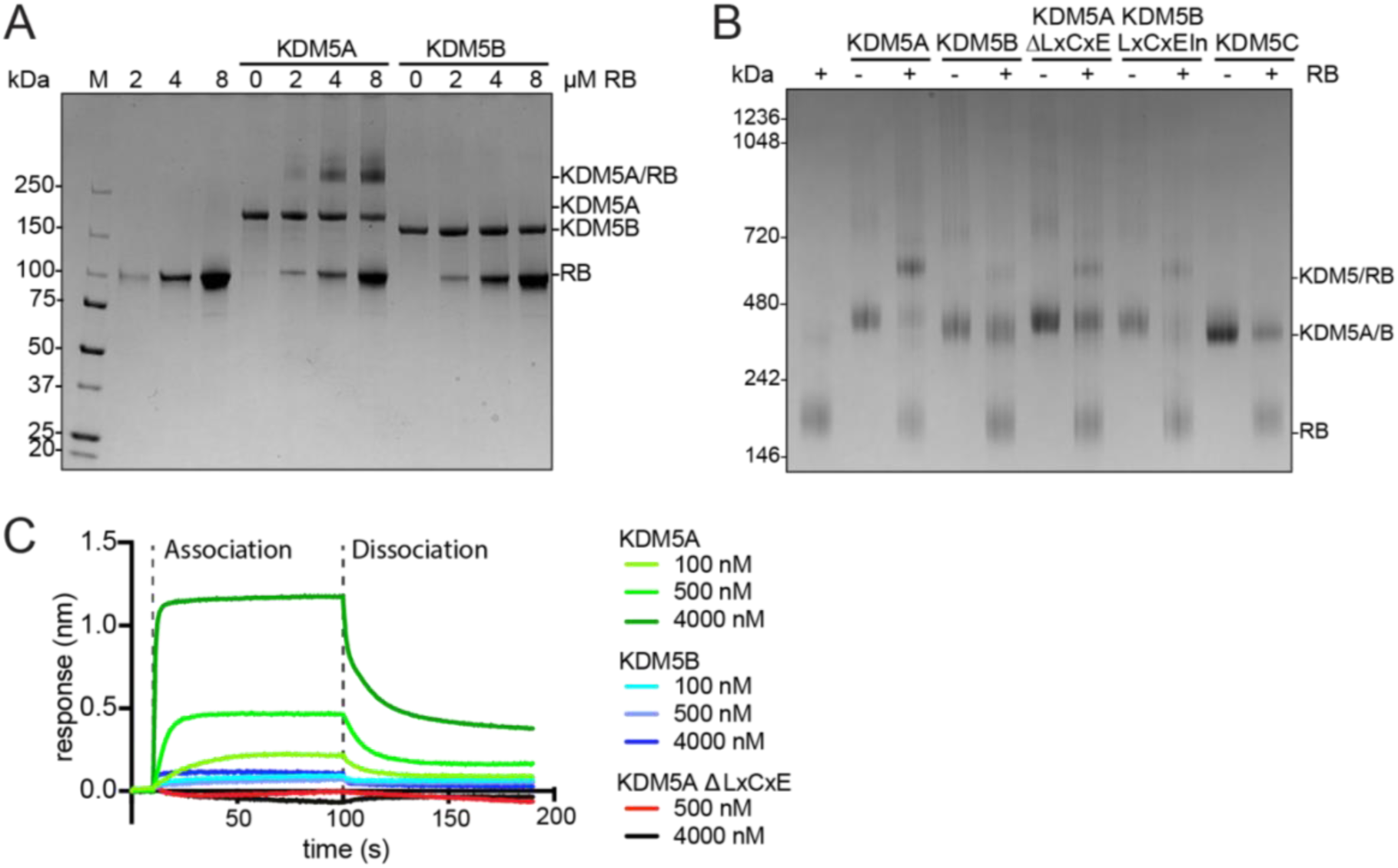
RB binding of KDM5A and KDM5B. (A) SDS-PAGE of 4 µM RB and 2 µM KDM5A or KDM5B cross-linked with Bis(sulfosuccinimidyl)suberate (BS^3^). (B) Blue native PAGE of 4 µM RB incubated with 2 µM WT KDM5A, WT KDM5B, KDM5A **Δ**LxCxE, KDM5B LxCxE insertion, or KDM5C. (C) Biolayer interferometry (BLI) sensogram of WT KDM5A (green), WT KDM5B (blue), or KDM5A **Δ**LxCxE (orange/red) binding to RB. Biotinylated RB was bound to the streptavidin chip, association and dissociation time points indicated by dashed lines (N = 3).

To test this hypothesis, we generated KDM5A constructs lacking the LxCxE motif (KDM5AΔLxCxE) and KDM5B construct where aa1366-aa1381 of KDM5A, including the LxCxE motif, was introduced into KDM5B at position 1314 (KDM5B LxCxE In). These constructs were also compared to KDM5C, which lacks both the LxCxE sequence and the NTE1A region. The loss of the LxCxE motif substantially reduced binding of KDM5A to RB as judged by native PAGE, while its introduction to KDM5B resulted in increased RB interaction. KDM5C did not show any interaction with RB (Fig. 2B). These observations were further supported by BLI experiments, in which clear binding of KDM5A to RB was detected (K_D_ of 136±32 nM), albeit with fast dissociation, indicating transient binding (Fig. 2C, Table 2). No RB binding to KDM5B was detectable with BLI, and deletion of the LxCxE motif from KDM5A abrogated its RB interaction (Fig. 2C). Taken together, these results suggest that the NTE1A region on its own is insufficient to mediate RB interactions *in vitro*.

**Table: 2.** Table of the binding constants obtained from biolayer interferometry. Means of at least 3 experiments and the Standard Error of the mean are shown.

| | KD (nM) | $k_{on}$ (1/Ms) | $k_{off}$ (1/s) |
| --- | --- | --- | --- |
| WT RB WT KDM5A | 136±32 | 2.25±0.81 x10<sup>5</sup> | 3.08±1.37 x10<sup>-2</sup> |
| RB LP WT KDM5A | 123±11 | 2.87±0.46 x10<sup>5</sup> | 3.49±0.36 x10<sup>-2</sup> |

To better define the interaction site between KDM5A and RB, we performed cross-linking mass spectrometry (XL-MS) with recombinant KDM5A and RB, using the chemical cross-linkers bis(sulfosuccinimidyl) suberate (BS^3^) and disuccinimidyl dibutyric urea (DSBU). Overall, 511 unique cross-links and 12 intermolecular cross-links were detected with the highest number of cross-linked residues located in solvent-exposed flexible linker regions, especially in KDM5A between the ARID and PHD1 domain and N-terminal of PHD3 and in an unstructured region C-terminal of the pocket B domain of RB (Fig. 3A). Published experimental structures were used to test whether the observed cross-links were in agreement with known experimentally determined structures of KDM5A and RB (Suppl. Fig. 7A,B). All of the 20 intra-molecularly cross-linked residues located within the KDM5A active core (PDB: 5CEH^[52]^) were within a distance of less than 30 Å between the Cα atoms, and only two exceeded a distance of 24 Å (Suppl. Fig. 7A). Of the 68 cross-links located in the RB-N and RB-LP domains (PDB: 4ELJ^[53]^), only one exceeded 30 Å Cα distance (Suppl. Fig. 7B).

**Figure 3:**
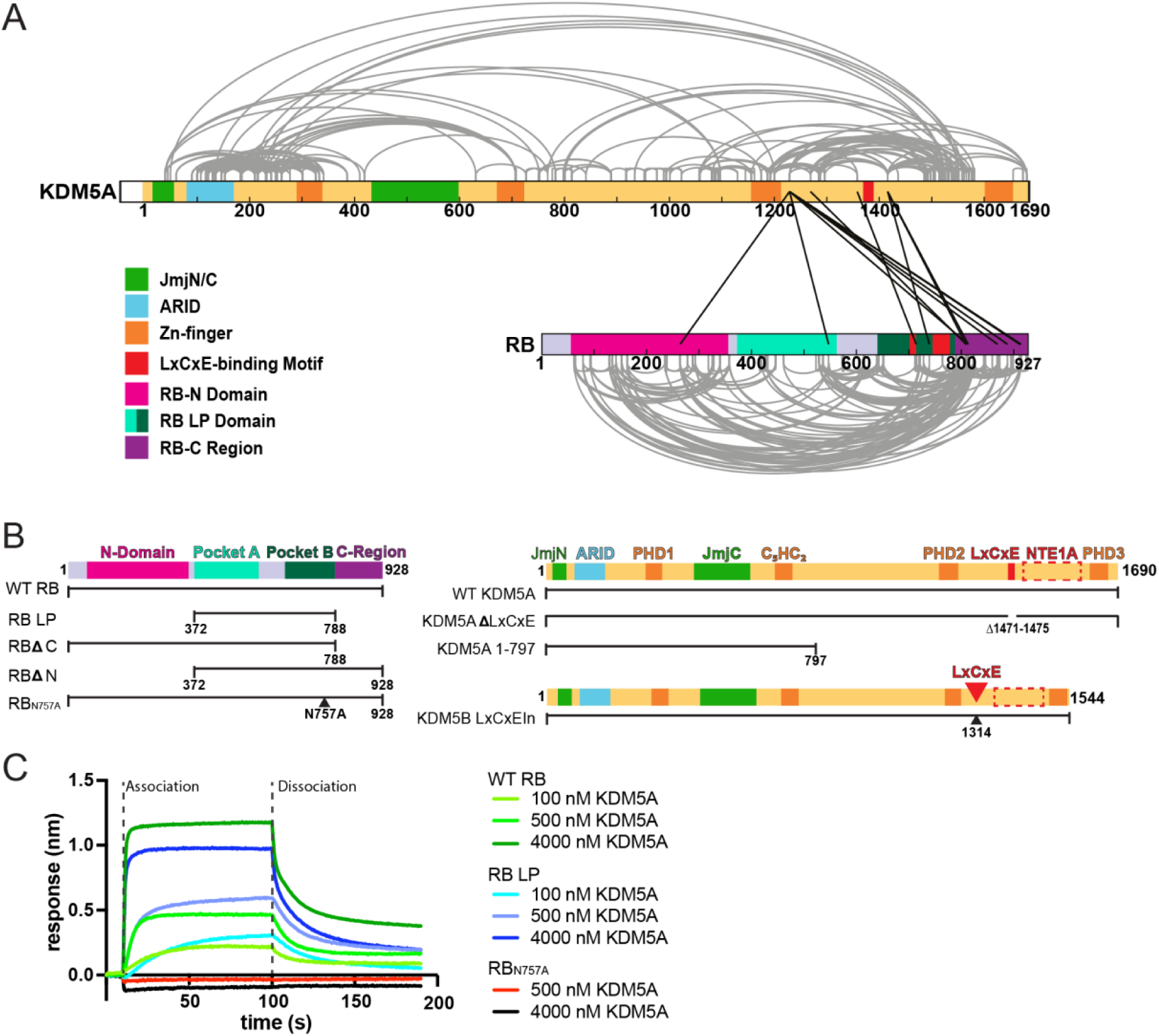
Determinants of RB binding by KDM5A (A) Overview of intra- and intermolecular cross-links found in KDM5A and RB with Bis(sulfosuccinimidyl)suberate (BS^3^) and disuccinimidyldibutyric urea (DSBU). (B) Schematic of constructs designed based on cross-linking results. Left: RB constructs: wildtype RB (WT RB), RB large pocket domain (RB LP), deletion of RB-C region (RBΔC), deletion of RB-N Domain (RBΔN), and point mutation of N757A in LxCxE binding site (RB_N757A_). Right, KDM5 constructs, including wild-type KDM5A (WT KDM5A), deletion of LxCxE sequence aa1471-1475 (KDM5A ΔLxCxE), KDM5A active core (KDM5A 1-797), and insertion of the KDM5A LxCxE motif in KDM5B (KDM5B LxCxE). (C) Biolayer interferometry (BLI) sensogram of WT KDM5A binding to either wild-type RB (WT RB) in green, RB large pocket domain (RB LP) in blue, or LxCxE binding mutant of RB (RB_N757A_) in orange/red. Biotinylated RB constructs were bound to the streptavidin chip; association and dissociation time points are indicated as dashed lines, and data represent three independent replicates (N=3).

Of the 12 intermolecular cross-links, four were found between the LxCxE motif in KDM5A and the respective LxCxE binding site in RB, while six cross-links were detected between a region C-terminal of PHD finger 2 of KDM5A to the C-terminal region of RB (Figure 3A). Our XL-MS results therefore confirm spatial proximity between the RB pocket domain and the LxCxE motif of KDM5A and also suggest a possible contribution of a region adjacent to PHD2 of KDM5A. To test whether regions other than the LxCxE motif contribute significantly to KDM5A/RB interactions, different RB constructs were generated, including deletions of either the RB-N domain (RBΔN) or the RB-C domain (RBΔC), and of the independent RB-N domain (RB-N), RB-LP domain (RB LP), RB pocket A (RB pA), and RB-C region (RB-C). Additionally, a reported non-binding mutant of the LxCxE binding site was generated by mutating glutamine 757 to an alanine (RB_N757A_) (Fig. 3B).^[54]^ A preliminary analysis by blue native PAGE showed that only RB constructs with an intact LxCxE binding pocket formed complexes with KDM5A (Suppl. Fig. 5C). These results were further verified by BLI experiments, in which the large pocket domain alone was sufficient to interact with KDM5A in the same manner as wild-type (WT) RB (K_D_ = 123±11 nM), while the N757A mutation led to a complete loss of interaction (Fig. 3C, Table 2). Therefore, we conclude that KDM5A, but not KDM5B, interacts stably with full-length RB *in vitro* and that this interaction is primarily mediated by LxCxE motif binding to its canonical binding site in the pocket domain of RB.

### KDM5A-RB interactions, nucleosome binding, and demethylase activity

Having established a biochemical system for studying the KDM5A-RB interaction, we characterized functional consequences on KDM5A activity. Since an impact of RB interactions on the demethylase activity of KDM5A has been hypothesized^[38,41]^, we performed a series of *in vitro* assays to test the interplay of both proteins in this recombinant context. First, we tested possible effects of RB on substrate nucleosome binding of KDM5A, using EMSAs with H3K4me3MLA nucleosomes. We found that RB can bind nucleosomes, with several RB molecules being able to interact with individual nucleosomes, as indicated by a shift of free nucleosomes to several slowly migrating species (Fig. 4A). Incubation of RB and KDM5A together with nucleosomes led to a new, distinct band, indicating the formation of a ternary KDM5A-RB-nucleosome complex. Since direct RB engagement with nucleosomes has not been described previously, we performed additional EMSA experiments to further characterize this binding. RB bound free double-stranded DNA with comparable efficiency to nucleosomes (Fig. 4B), while binding was reduced using nucleosomes without linker DNA (Fig. 4C). Together, these results indicate that RB-nucleosome interactions are driven by DNA binding rather than histone protein interactions. To narrow down the RB regions required for nucleosome binding, we mapped the RB domains using various deletion constructs. Here, we observed that only constructs containing the RB-C region formed distinct nucleosome bands, whereas constructs lacking the RB-C region showed reduced binding (Suppl. Fig. 7D). We further examined the effects of RB association on KDM5A demethylase activity. When H3_1-18_K4me3 peptides were used as a substrate, no significant effect on the reaction rate or the substrate affinity was observed (Suppl. Fig. 8A,B, Table 3). Similarly, H3K4me3MLA nucleosomes were demethylated by KDM5A as efficiently in the presence as in the absence of RB (Fig. 4D). These results show that RB can bind to nucleosomes primarily through interactions between the RB-C region and nucleosomal DNA. As the RB-C region is rich in basic aa (Suppl. Fig. 8C), electrostatic interactions with the phosphate backbone of the DNA are conceivable. RB binding did not strongly affect nucleosome binding or the demethylase activity of KDM5A using either peptide or nucleosome substrates and under the conditions tested. This indicates that KDM5A can efficiently bind and demethylate nucleosomes in the presence of RB, suggesting that the functional consequences of RB interactions are likely mediated by alternative mechanisms.

**Figure 4:**
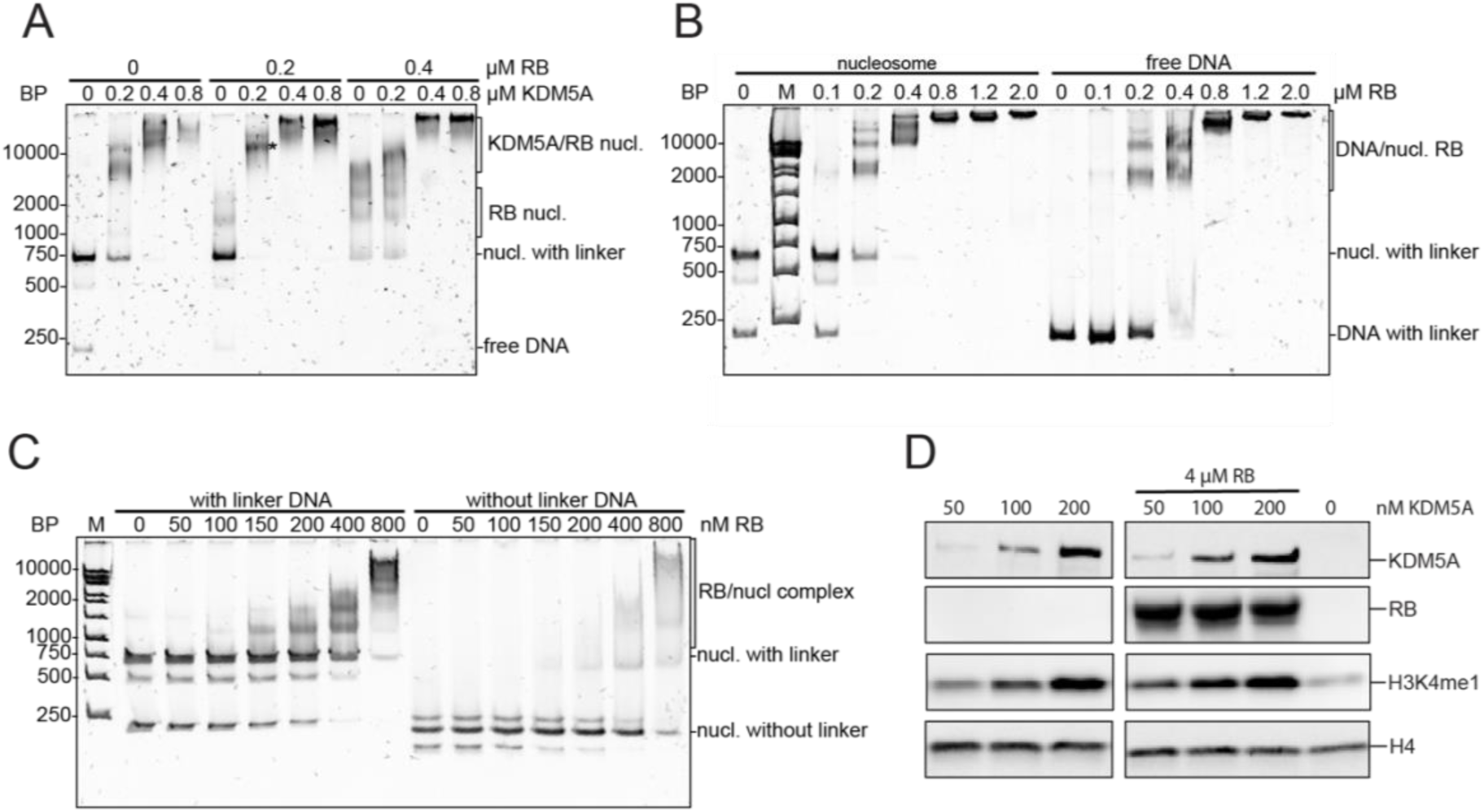
Implications of RB/KDM5A interaction (A) Electrophoretic mobility shift assays (EMSAs) of KDM5A and RB with 100 nM of H3K4me3MLA nucleosomes (nucl.) with linker DNA. The band corresponding to the putative ternary complex of KDM5A, RB, and nucleosome is indicated with an asterisk (*). (B) EMSAs of RB with H3K4me3MLA nucleosomes (left lanes) or DNA alone (right lanes) at 100 nM concentration each. (C) EMSA of RB with 100 nM H3K4me3MLA nucleosomes with and without linker DNA. Complex formation was analyzed by non-denaturing 5% polyacrylamide gel electrophoresis with SYBR Gold staining. (D) Western blot of KDM5A demethylase activity on H3K4me3MLA nucleosome with linker DNA in the presence or absence of RB (N=3).

**Table: 3.** Table showing the k_cat_ and K_M_ of KDM5A with different concentrations of RB added, obtained from FDH coupled assays with H3_1-18_K4me3 peptides. Means of at least three experiments and the Standard Error of the mean are shown.

| | 0 μM RB | 2 μM RB | 4 μM RB | 8 μM RB |
| --- | --- | --- | --- | --- |
| $k_{cat}$ (1/min) | 0.37±0.11 | 0.37±0.10 | 0.31±0.11 | 0.34±0.12 |
| $K_M$ (μM) | 144±77 | 126±69 | 123±64 | 107±38 |

## Discussion

While KDM5A and KDM5B are similar in both structural and functional aspects, and it has been reported that different KDM5 enzymes can rescue each other’s loss to some extent^[20]^, both perform essential and non-redundant roles.^[26–29]^ To biochemically characterize the differences between the two enzymes, we systematically compared key aspects of KDM5A and B functions *in vitro*. We identified differences between full-length KDM5A and KDM5B regarding their demethylase activities, nucleosome binding, and interactions with RB.

Our negative staining results, combined with rigid-body fitting of AlphaFold3 predictions, underscore the similar overall form and architecture of KDM5A and KDM5B. Both featured a compact N-terminal core comprising the JmjN, JmjC, ARID, PHD1, and the C5HC2-type zinc finger domains, from which the C-terminal part extends in an elongated structure including PHD2/3 and additional interaction sites such as the LxCxE sequence in KDM5A and the NTE1A region in both. For KDM5B, the observations agree with a previous study.^[50]^ To identify more detailed structural differences between KDM5A and B, high-resolution investigations will be required.

Using H3_1-18_K4me3 peptides as substrates, we observed differences in the enzyme kinetics of KDM5A and KDM5B, with a 20-fold higher substrate affinity of KDM5B over that of KDM5A, but a higher conversion rate by KDM5A at high substrate concentrations, using identical conditions for both enzymes. The demethylase activities we determined for full-length KDM5A and B show relatively slow reaction rates (k_cat_) but similar substrate affinities (K_M_) compared to other investigations that focused on substrate-specific Michaelis-Menten kinetics using FDH-coupled assays with H3K4me3 peptides and truncated versions of KDM5 enzymes. For KDM5A 1-797, a higher conversion rate with a k_cat_ of 2,79±0.05 min^-1^ and a K_M_ of 58±5 µM^[14]^ was reported, compared to our results with a k_cat_ of 0.466±0.105 min^-1^ and a K_M_ of 106.9±49.7 µM (Fig. 1E-G, Table 1). Similar results were reported for truncated KDM5B 1- 755 with a conversion rate k_cat_ of 2.00±0.03 min^-1^ and a K_M_ of 4.5±0.5 µM^[13]^ compared to our results with a k_cat_ of 0.303±0.075 min^-1^ and a K_M_ of 5.3±2.2 µM for KDM5B (Fig. 1E-G, Table 1).^[13,50,55]^. We did not observe differences between the activities of full-length versus truncated KDM5A and B variants, so the slower rates we observe relative to published values likely reflect differences in reaction conditions.

KDM5A showed stronger substrate nucleosome binding than KDM5B, which is in seeming contradiction to the kinetic parameters determined in peptide demethylation assays. These observations underscore that, while the demethylation of isolated histone peptides informs on fundamental enzyme kinetics, nucleosome binding is likely to affect the demethylation of this more complex, physiological substrate. Investigations of other JmjC demethylases have shown that close homologs can exhibit important differences in nucleosome binding and demethylation. For example, the nucleosome binding of human KDM2A and KDM2B that target H3K36me2 shows key differences. KDM2A but not KDM2B has an IDR that binds to the H2A/H2B acidic patch, and this interaction was proposed to help position KDM2A for partial DNA unwrapping to aid H3K36me2 demethylation.^[56]^ Interestingly, an impact of the KDM5A IDR, which lies between the PHD1 zinc finger and the ARID domain, on nucleosome binding and demethylation has been shown^[49]^, suggesting that this IDR, similar to KDM2, aids in the correct positioning of KDM5A towards H3K4me3. Of the three arginine-rich patches of KDM5A that were proposed to mediate acidic patch interactions, KDM5B lacks one. It is conceivable that KDM5A binds the nucleosome more stably due to this interaction and that KDM5B interacts only transiently because this interaction is missing. Future studies will address the mechanistic basis of nucleosome interactions of KDM5A and B, and how these interactions affect chromatin association and histone demethylation in the context of other interaction partners and in the cell.

The interactions with other regulatory factors are a key to chromatin regulator function.^[1,2,57]^ The first reported KDM5 interactor is RB, which was shown to associate with KDM5A and B even before their demethylase activities were characterized.^[30,42,58]^ Despite this long-known relationship, the biochemical basis and implications of the KDM5A RB interplay have remained poorly understood. Therefore, we systematically studied the interactions between KDM5A and KDM5B with RB *in vitro* using recombinant human proteins. We found that RB and KDM5A formed a stable complex mediated by the LxCxE motif of KDM5A interacting with the canonical binding site in the pocket domain of RB. Deletion of the LxCxE motif or inactivation of the LxCxE binding pocket lead to a loss of KDM5A-RB interaction. Accordingly, KDM5B, which lacks the LxCxE motif, did not stably interact with RB unless the KDM5A LxCxE motif was artificially introduced. The NTE1A region has been proposed as an additional interaction interface between KDM5A and KDM5B with RB. This binding mode was detected in cellular systems upon ectopic overexpression of truncation constructs followed by immunoprecipitation (IP).^[42,46,47,59]^ Since we could not detect measurable binding between KDM5 proteins and RB in the absence of the LxCxE motif, we propose that the NTE1A region may contribute only marginally to the interaction. Alternatively, the association detected by co-IP was mediated by additional interaction partners that may co-precipitate with KDM5B or RB in this setup. For example, other KDM5B interactors, such as SIN3 or NuRD co-repressor complexes^[8,22,60]^, may in turn interact with RB^[61,62]^ and co-precipitate with KDM5B.

One possible functional implication of the proposed RB-KDM5 interaction is an effect of RB binding on KDM5 activity in cells.^[29,41]^ To test whether RB binding directly affects KDM5A activity, we performed demethylase activity assays in the presence or absence of RB. Under the conditions tested here, there was no strong effect of RB binding on histone peptide or nucleosome demethylation by KDM5A, and we therefore conclude that RB is unlikely to regulate KDM5A function through direct modulation of its enzymatic activity.^[29,40,63]^ We showed that nucleosome binding and demethylation are not negatively affected by RB *in vitro*. In the cellular context, it is still possible that RB binding may affect KDM5A recruitment or chromatin dwell times, for example, in the context of E2F target genes.^[37]^ Furthermore, KDM5A-RB interactions may impact the targeting, formation, and stability of chromatin regulator complexes that include either one or both of the proteins. Such effects would not be captured by our recombinant system and require more detailed investigations in cellular experimental systems.

We observed that RB binds directly to nucleosomes in the absence of additional interaction partners and showed that this interaction is mainly driven by DNA binding of the C-terminal region of RB, which is rich in basic amino acids and includes many phosphorylation sites.^[34]^ It will be a valuable subject of future studies to investigate whether direct chromatin binding by RB affects its cellular function in addition to its interactions with KDM5A, transcription factors such as E2F, and other chromatin regulators.

Finally, it will also be important to determine the extent to which KDM5A binding affects RB function. For example, binding of KDM5A to the RB pocket domain may compete with other pocket-domain interactors and thus antagonize their function. Additionally, KDM5A interactions may have an impact on RB phosphorylation. For instance, it has been proposed that the phosphorylation of RB may be regulated by KDM5 interactions in melanoma cells.^[42]^ Vice versa, RB phosphorylation was shown to regulate LxCxE-dependent binding to RB^[64]^, raising the possibility that KDM5A interactions may be affected by the phosphorylation state of RB. Therefore, more detailed, mechanistic, and cellular studies will have to be performed to decipher the basis and implications of the interplay of these critical chromatin regulators in various physiological and disease contexts.

Collectively, our findings establish a biochemical framework for studying the distinct properties of KDM5A and KDM5B, providing a foundation for future studies to investigate how these demethylases function distinctly within broader chromatin regulatory networks. The mechanistic distinction in RB binding, which primarily depends on the LxCxE motif unique to KDM5A, may help to explain why these paralogs have non-redundant roles in cell cycle regulation and cancer.

## Supporting information

Cross-links BS3 InSolution

Cross-links DSBU InGel

Cross-links BS3 InGel

Cross-links BS3 InGel

Sample_Legend_and_LC-MS_Settings_v01.docx.

## Acknowledgements

We thank Uli Baumann and Jan Gebauer for their support in protein interaction methods and access to the protein interaction infrastructure of the PIP_C_, Monika Gunkel and Elmar Behrmann for their support regarding cryo-EM methods. We acknowledge access to the cryo-EM infrastructure of StruBiTEM (Cologne, funded by DFG Grant 357484240). We would like to thank Jenny Bormann for her excellent technical support.

## Author contributions

Conceptualization and experiment design by T.R. and S.P.. Experimental work performed by T.R., J.R., E.P., F.B., M.D., F.K., and A.H.. Data processing, analysis and interpretation by T.R., E.P., F.B., F.K., L.Z., B.S., A.R., and M.K.. T.R. and S.P. prepared the manuscript with input from all coauthors.

## Data and materials availability

Negative-stain EM maps are available on EMDB: (…).^[65]^

Cross-linking MS Data are uploaded and available on Proteomics Identification Database (PRIDE)^[66]^ under PRIDE ID: PXD075614

## Methods

### Plasmids and antibodies

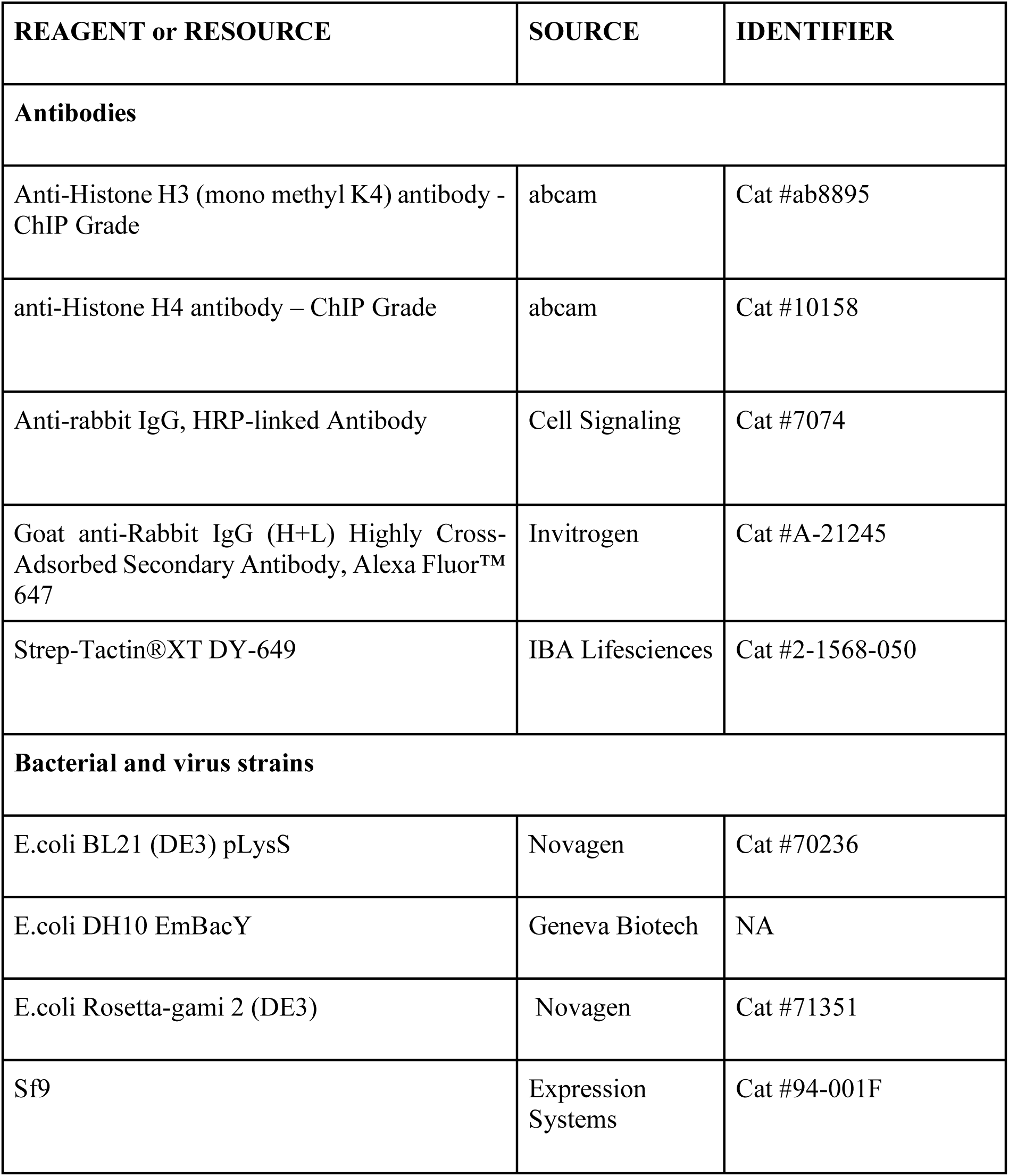
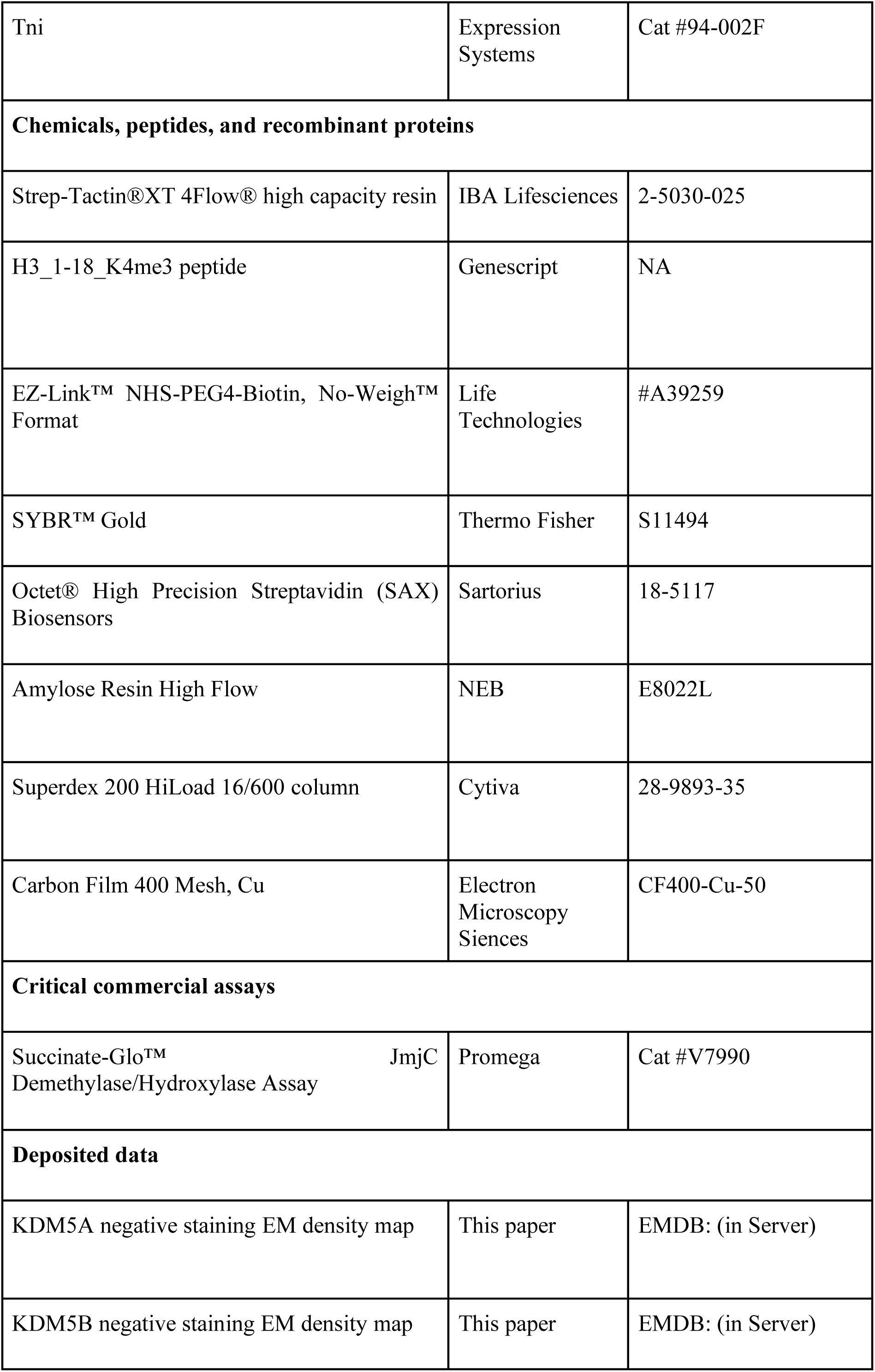
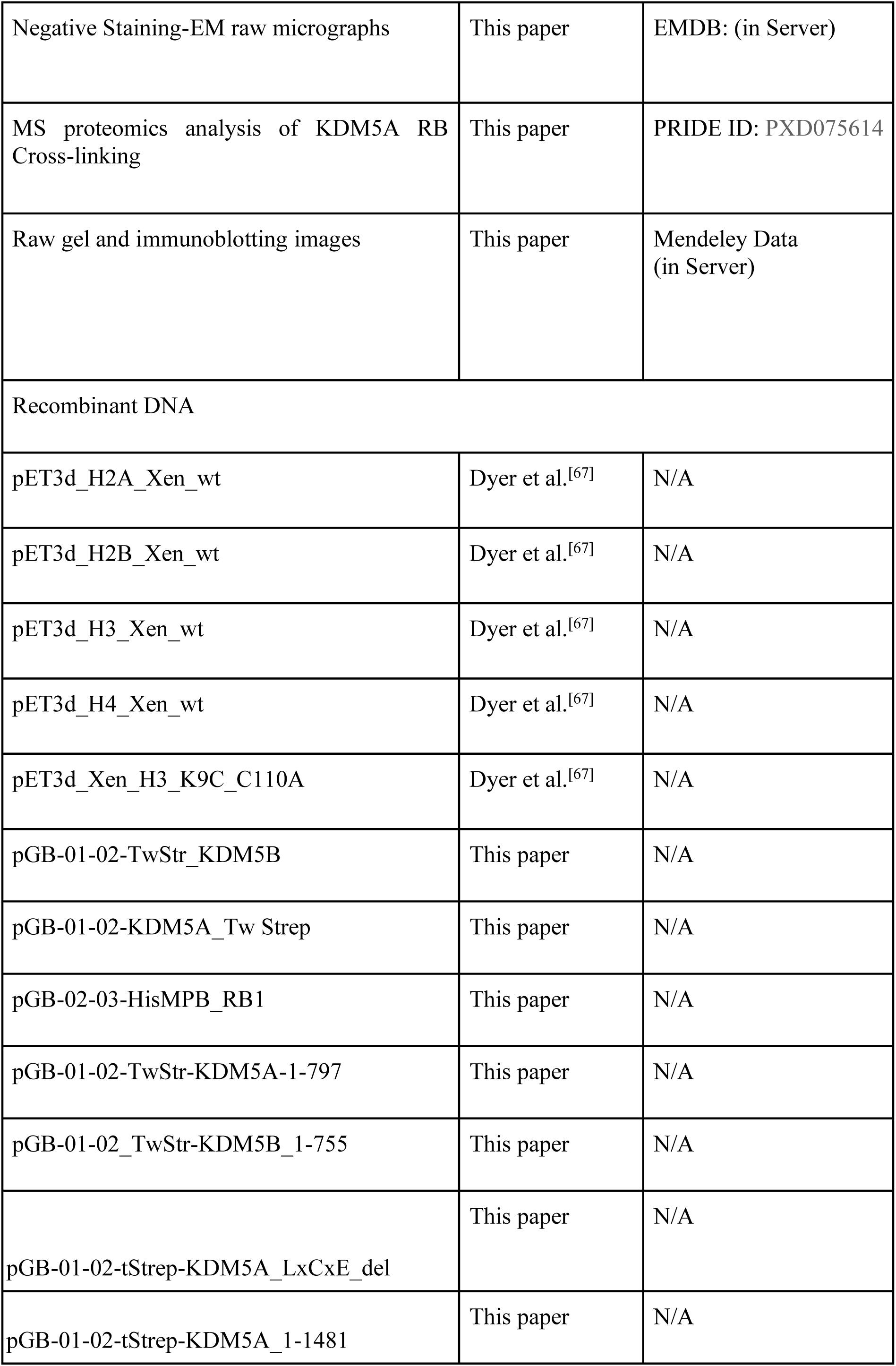
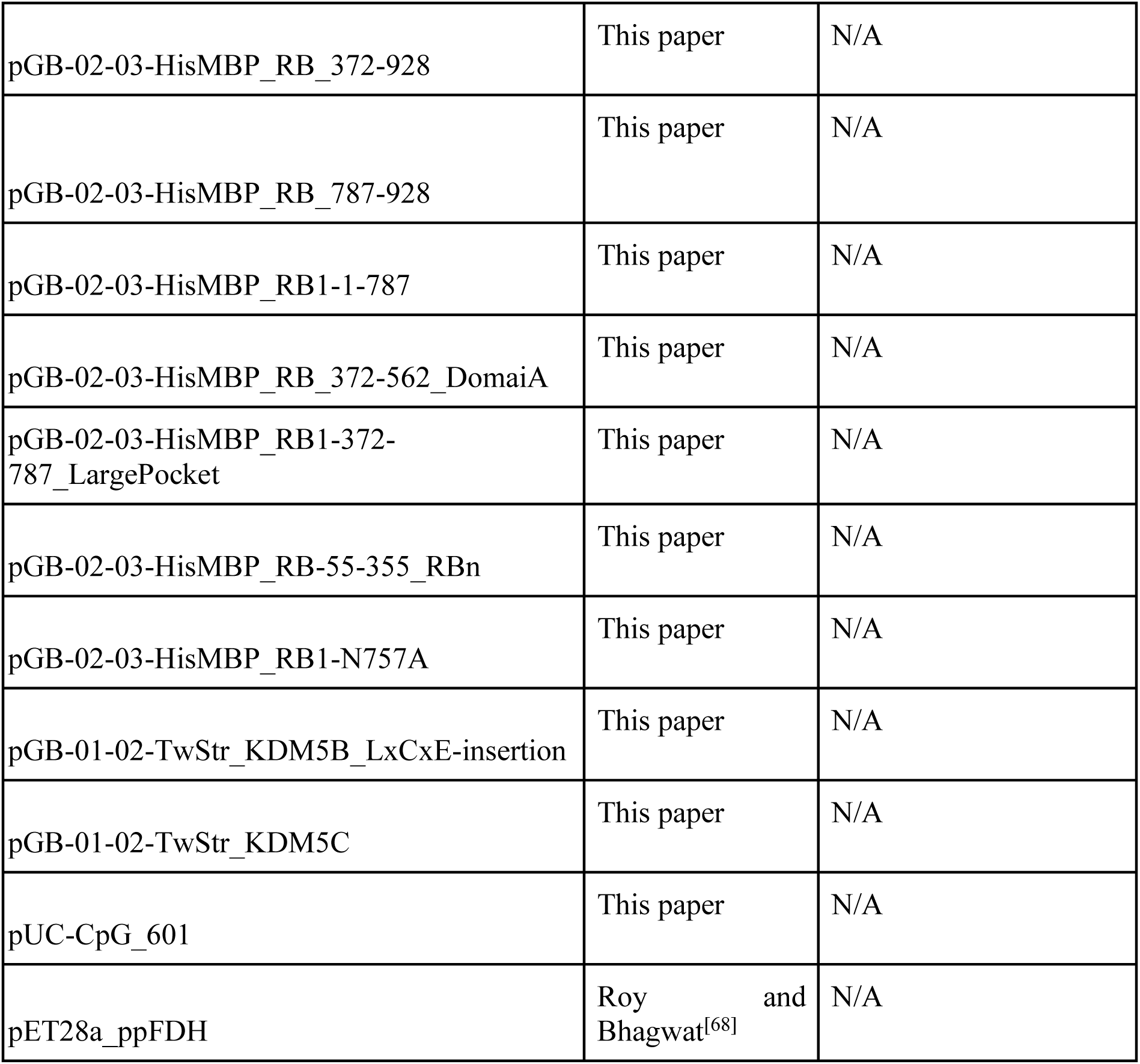

## Methods

### KDM5 constructs cloning, expression, and purification

Codon-optimized coding sequences of KDM5A/B (N-terminally StrepII-tagged) were purchased from Twist Bioscience and cloned into GoldenBac vectors (pGB^[69]^) for insect cell expression. DNA fragments coding for the indicated KDM5 constructs were generated by PCR and inserted into GoldenBac vectors (pGB^[69]^) using the In-Fusion® Snap Assembly Master Mix (Takara Bio).

Vectors were introduced into E. coli DH10 EmBacY (Geneva Biotech) by heat shock. Baculoviral stocks were generated in Sf9 cells (Expression Systems) cultivated at a starting density of 1 x 10^6^ cells/well in coated 6-well tissue culture dishes at 28 °C using protein-free ESF 921 insect cell culture medium (Expression Systems) supplemented with penicillin/streptomycin (Life Technologies) until fluorescence was observed (72-120 h). Cells were pelleted, and the supernatant was either stored at 4 °C or used directly for virus amplification. For virus amplification, 50 ml of Sf9 cells (Expression Systems) with a density of 1 x 10^6^ cells in protein-free ESF 921 insect cell culture medium (Expression Systems) supplemented with penicillin/streptomycin (Life Technologies) were infected with the baculovirus and incubated at 28 °C for 96 h. For protein expression, *Trichoplusia ni* cells (Tni, Expression Systems) were infected with 2 %v/v baculovirus at a density of 1 x 10^6^ cells at 28 °C in a shaker incubator using ESF 921 Insect Cell Culture Medium (Expression Systems) supplemented with penicillin/streptomycin (Life Technologies) and 0.5 mM ZnCl_2_. Cells were shaken at 28 °C for 72 h, then harvested and stored at −80 °C until further use.

Insect cell pellets were thawed and resuspended in cold lysis buffer (50 mM KPO_4,_ pH 7.7, 300 mM NaCl, 1 mM DTT, 5 % glycerol, 1.5 mM MgCl_2_, 0.1 % NP-40, 1 µM Pepstatin, 10 µM Bestatin, 2 µg/ml Aprotinin, 10 µM E64, 10 µM Leupeptin, 5 U/ml *S. marcescens* nuclease^[70]^, cOmplete™ mini EDTA-free protease inhibitors (Roche)). The suspension was lysed by sonication (15 s on, 45 s off, 4 min on time, 50 % amplitude), and centrifuged at 23000 xg for 30 min. The supernatant was incubated with Strep-Tactin®XT 4Flow® high capacity resin (IBA Lifesciences) for 1 h at 4° C, applied to a gravity flow column, washed with 5 column volumes (CV) lysis buffer, followed by 5 CV wash buffer (50 mM KPO_4_, pH 7.7, 600 mM NaCl, 1 mM DTT, 2 % glycerol) and 5 CV 50 mM KPO_4_, pH 7.7, 600 mM NaCl, 1 mM DTT, 2 % glycerol before elution using 50 mM KPO_4_, pH7.7, 100 mM NaCl, 1 mM DTT, 2 % glycerol supplemented with 50 mM biotin. The eluted fractions were concentrated and further purified by SEC on a Superdex 200 HiLoad 16/600 column (Cytiva) in a buffer containing 50 mM KPO_4_, pH 7.7, 100 mM NaCl, 1 mM DTT, and 2% glycerol. Peak fractions were pooled, concentrated, and used for further experiments, or flash-frozen in liquid nitrogen and stored at −80 °C. For the BLI assay, the StrepII tags were removed by overnight incubation with TEV protease at 4 °C before the SEC purification step.^[71]^

### RB constructs cloning, expression and purification

RB 413 pSG5L HA RB was a gift from William Sellers (Addgene plasmid # 10720; http://n2t.net/addgene:10720; RRID: Addgene_10720).^[71]^ The HA-tag was replaced with a His-MBP tag (N-terminal His-MBP-tagging) and cloned into GoldenBac vectors (pGB^[69]^) for insect cell expression. DNA fragments coding for the indicated RB constructs were generated by PCR and inserted into GoldenBac vectors (pGB^[69]^) using the In-Fusion® Snap Assembly Master Mix (TakaraBio).

Vectors were introduced into E. coli DH10 EmBacY (Geneva Biotech) by heat shock. Baculoviral stocks were generated in Sf9 cells (Expression Systems) cultivated at a starting density of 1 x 10^6^ cells/well in coated 6-well tissue culture dishes at 28 °C using protein-free ESF 921 insect cell culture medium (Expression Systems) supplemented with penicillin/streptomycin (Life Technologies) until fluorescence was observed (72-120 h). Cells were pelleted, and the supernatant was either stored at 4 °C or used directly for virus amplification. For virus amplification, 50 ml of Sf9 cells (Expression Systems) with a density of 1 x 10^6^ cells in protein-free ESF 921 insect cell culture medium (Expression Systems) supplemented with penicillin/streptomycin (Life Technologies) were infected with the baculovirus and incubated at 28 °C for 96 h. For protein expression, *Trichoplusia ni* cells (Tni, Expression Systems) were infected with 2 %v/v baculovirus at a density of 1 x 10^6^ cells at 28 °C in a shaker incubator using ESF 921 Insect Cell Culture Medium (Expression Systems) supplemented with penicillin/streptomycin (Life Technologies). Cells were shaken at 28 °C for 72 h, then harvested and stored at −80 °C until further use.

Insect cell pellets were thawed and resuspended in cold lysis buffer (50 mM KPO_4,_ pH 7.7, 300 mM NaCl, 1 mM DTT, 5 % glycerol, 1.5 mM MgCl_2_, 0.1 % NP-40, 1 µM Pepstatin, 10 µM Bestatin, 2 µg/ml Aprotinin, 10 µM E64, 10 µM Leupeptin, 5 U/ml *S. marcescens* nuclease^[70]^, cOmplete™ mini EDTA-free protease inhibitors (Roche)). The suspension was lysed by sonication (15 s on, 45 s off, 4 min on time, 50 % amplitude), and centrifuged at 23000 xg for 30 min. The supernatant was incubated with Amylose Resin High Flow (NEB) for 2.5 h at 4° C then applied to a gravity flow column, washed with 5 CV lysis buffer, followed by 5 CV wash buffer (50 mM KPO_4_, pH 7.7, 600 mM NaCl, 1 mM DTT, 2 % glycerol) and 5 CV 50 mM KPO_4_, pH 7.7, 600 mM NaCl, 1 mM DTT, 2 % glycerol before elution using 50 mM KPO_4_, pH7.7, 100 mM NaCl, 1 mM DTT, 2 % glycerol supplemented with 50 mM maltose. The MBP-tag was removed by overnight incubation with TEV protease at 4 °C.^[71]^ The eluted fractions were concentrated and further purified by SEC on a Superdex 200 HiLoad 16/600 column (Cytiva) in a buffer containing 50 mM KPO_4_, pH 7.7, 100 mM NaCl, 1 mM DTT, and 2% glycerol. Peak fractions were pooled, concentrated, and used for further experiments, or flash-frozen in liquid nitrogen and stored at −80 °C.

### Purification of formaldehyde dehydrogenase (FDH)

The plasmid encoding *Pseudomonas putida* formaldehyde dehydrogenase (pET28a_ppFDH) was a kind gift from Prof. Ashok S. Bhagwat, Wayne State University. ^[68]^ FDH was expressed and purified as described previously.^[68]^ Briefly, pET28a_ppFDH was transformed into chemically competent *E.coli* Rosetta-gami 2 (DE3) (Novagen) by heat shock. Bacteria were grown in LB-broth (Sigma-Aldrich) supplemented with 50 µg/ml kanamycin. Expression was induced by the addition of 0.5 mM IPTG at an OD_600_ of 0.4 and expressed at 800 RPM at 25 °C overnight. then harvested and stored at −80 °C until further use. Bacterial pellets were resuspended in cold buffer 50 mM KPO_4_, pH 7.7, 100 mM NaCl, 25 mM imidazole, 1 µM Pepstatin, 10 µM Bestatin, 2 µg/ml Aprotinin, 10 µM E64, 10 µM Leupeptin, 5 U/ml *S. marcescens* nuclease^[70]^, cOmplete^TM^ mini EDTA-free protease inhibitors (Roche). The resuspension was lysed by sonication (15 s on, 45 s off, 4 min on time 50 % amplitude), and centrifuged at 2,300 x g for 30 min. The supernatant was loaded on a HiTrap chelating HP column (Cytiva) with a Fast Protein Liquid Chromatography System (Bio-Rad) loaded with Zn^2+^. The column was washed with 5 CV of 50 mM KPO_4_, pH 7.7, 100 mM NaCl, and 25 mM imidazole, then eluted with an imidazole gradient to 250 mM. The eluted fractions were concentrated. The protein was further purified by SEC on a Superdex S200 increase 10/300 column (Cytiva) in buffer containing 50 mM KPO_4_, pH 7.7, 100 mM NaCl, 1 mM DTT, 2 % glycerol. Peak fractions were pooled, concentrated to 5-10 mg/ml, and flash frozen in liquid nitrogen and stored at −80 °C.

### Nucleosome reconstitution

Histone proteins H2A, H2B, H3, and H4 were expressed and purified as described. ^[72]^ To later introduce methyl-lysine analogs (MLA)^[51]^, an H3 K4C C110A mutant was expressed. The trimethyl-MLA was then chemically introduced using (2-Bromethyl)-trimethylammoniumbromid as described.^[51]^ Two nucleosomal DNA constructs were used: either only the Widom ‘601’ nucleosome positioning sequence^[73]^ or the same sequence flanked by 40 bp of linker DNA at both ends (CACGCGACTGTGTGCCCGTCAGACGCTGCGCTGCCGGCGG<u>CTGGAGAATCCCGG</u> <u>TGCCGAGGCCGCTCAATTGGTCGTAGACAGCTCTAGCACCGCTTAAACGCACGTA</u> <u>CGCGCTGTCCCCCGCGTTTTAACCGCCAAGGGgATTACTCCCTAGTCTCCAGGCAC</u> <u>GTGTCAGATATATACATCCTGT</u>ATGCATGCATATCATTCGATCGGAGCTCCCGATC GATGC Widom 601 sequence underlined). The DNA was amplified by PCR, purified on an HiTrap® Q High Performance (Cytiva), and further purified by ethanol precipitation.^[74]^ For nucleosome assembly, equimolar amounts of all histones in 6 M GuHCl, 20 mM Tris/HCl, pH 7.5, 5 mM DTT were dialyzed into 2 M NaCl, 10 mM Tris/HCl, 5 mM EDTA, and the octamer was purified using a Superdex 200 increase 10/300 size exclusion column (Cytiva). The DNA and octamer were mixed at a 1:1 ratio and purified using a Bio-Rad 491 Prep-Cell after overnight salt-gradient dialysis, as described previously.^[67]^

### Formaldehyde dehydrogenase (FDH) coupled demethylase assay

For an FDH coupled KDM5 demethylase assay, 2 µM enzyme was incubated for 10 minutes at RT with 100 µM α-ketoglutarate, 10 µM (NH_4_)_2_Fe(SO_4_)_2_, 40 µM ascorbic acid, 0.5 µM FDH, 200 µM NAD^+^ in 50 mM KPO_4_, pH 7.7, 100 mM NaCl, 0.4 % glycerol carry-over. The reaction was started by adding 0.1-600 µM histone H3_1-18_K4me3 peptide (Genscript) directly before measurement. The formation of NADH was monitored by excitation at 280 nm and emission at 480 nm using a Hidex Sense Microplate Reader. The reaction was performed at 25 °C. Additionally, a NADH standard and negative controls without substrate and/or enzyme were measured. For each condition, at least two technical replicates were performed.

To calculate Michaelis-Menten kinetics, product formation was determined using the NADH standard row in Microsoft Excel. Due to the nonspecific activity of FDH against glycerol, the background activity was subtracted using the negative controls lacking KDM5s. Initial molecular reaction rates were calculated by linear regression and converted into reaction rates by dividing by the enzyme concentrations. The reaction rates were then plotted against substrate concentration, and K_M_ and k_cat_ were calculated directly using the Michaelis-Menten nonlinear regression from GraphPad Prism 11.0.1. Each experimental condition was repeated at least three times independently, and the average Michaelis-Menten kinetics with their standard error of the mean were calculated in GraphPad Prism 11.0.1. Significance between conditions was calculated in GraphPad Prism 11.0.1 using a one-way ANOVA, with P<0.05 considered statistically significant.

### Luminescence-based demethylase assay

Succinate formation was assessed using the Succinate-Glo™ JmjC Demethylase/Hydroxylase Assay from Promega. Therefore, 2 µM enzyme was incubated for 10 minutes at RT with 100 µM α-ketoglutarate, 10 µM (NH_4_)_2_Fe(SO_4_)_2_, 40 µM ascorbic acid, 0.5 µM FDH, 200 µM NAD^+^, in 50 mM KPO_4_, pH 7.7, 100 mM NaCl, and 0.4 % Glycerol carryover. The reaction was initiated by adding the 0.1-600 µM H3_1-18_K4me3 peptide, incubated for 4 min at 25 °C, and then stopped by incubation at 98 °C for 2 min. Afterward, the reaction was cooled to 25 °C, and 10 µl of the mixture was added to 10 µl of Succinate Detection Reagent I in a 384-well plate, then incubated at RT for 1 h. 20 µl of Succinate Detection Reagent 2 was added, and the mixture was incubated at RT for at least 10 min. Succinate was quantified by luminescence using a Hidex Sense Microplate Reader. For each condition, at least three technical replicates were performed, along with a background measurement without enzyme and a standard row with succinate in the reaction buffer.

To calculate Michaelis-Menten kinetics, product formation was determined using the Succinate standard row using Microsoft Excel. Background activity was subtracted using the negative controls lacking KDM5 enzymes. Initial molecular reaction rates were calculated by dividing by the reaction time of 2 min and converted into reaction rates by dividing by the enzyme concentrations. The reaction rates were then plotted against substrate concentration, and K_M_ and k_cat_ were calculated directly using the Michaelis-Menten nonlinear regression from GraphPad Prism 11.0.1. Each experimental condition was repeated at least three times, and the average Michaelis-Menten kinetics with their standard error of the mean were calculated in GraphPad Prism 11.0.1. Significance between conditions was calculated in GraphPad Prism 11.0.1 using a one-way ANOVA, with P<0.05 considered significant.

### Western Blot activity assays

For the Western blot-based activity assays, either 100, 200, or 400 nM KDM5A was mixed with a final concentration of 4 µM RB in 10 µM (NH_4_)_2_Fe(SO_4_)_2_, 40 µM ascorbic acid, 0.5 µM FDH, 200 µM NAD^+^ in 50 mM KPO_4_, pH 7.7, 100 mM NaCl, and incubated for 30 min on ice. The reaction was initiated by adding 150 nM H3K4me3MLA nucleosome and 1 mM α-ketoglutarate. The reaction mixture was incubated at 23 °C for 2 min, then stopped by the addition of 5x SDS buffer and heated to 98 °C.

Samples were applied to Mini-PROTEAN TGX Stain-Free Precast 4–20 % gradient gels and run at 110 V for 90 min. Subsequently, gels were blotted onto 0.2 µm PVDF and blocked with 5 % BSA in TBS-T. Respective antibodies were diluted as indicated, Strep-Tactin®XT DY-649 (#2-1568-050, IBA Lifesciences) 1:2000 for KDM5s, Anti-Histone H3 (mono methyl K4) antibody - ChIP Grade (ab8895, abcam) 1:1000, and Anti-Histone H4 antibody - ChIP Grade (ab10158, abcam) 1:1000, in TBS-T were applied o/n at 4 °C. Membranes were washed three times with TBS-T, and secondary antibodies, anti-rabbit IgG HRP-linked (#7054, Cell Signaling) diluted 1:2500 in TBS-T for H3K4me1 and goat anti-rabbit IgG (H+L) Highly Cross-Adsorbed Secondary Antibody, Alexa Fluor™ 647 (A-21245, Invitrogen) for H4 diluted 1:5000 in TBS-T were applied at RT for 1 h. Membranes were washed and either dried or used for imaging with ECL Prime Reagent (Cytiva).

### Blue-native PAGE

2 µM of KDM5 WT or mutants in reaction buffer (50 mM KPO_4_, pH 7.7, 100 mM NaCl) were mixed with 4 µM RB WT or mutants and incubated on ice for at least 10 min. Immediately before loading to a 3-12 % gradient NativePAGE™ Bis-Tris Mini Protein Gels (Invitrogen), samples were mixed with NativePAGE™ sample buffer (4X)(Invitrogen). NativePAGE™ running buffer (Invitrogen) was used, and NativePAGE™ cathode buffer additive (Invitrogen) was added to the cathode buffer.

### Electrophoretic Mobility Shift Assay (EMSA)

100 nM nucleosomes in reaction buffer (50 mM KPO_4_, pH 7.7, 100 mM NaCl) were mixed with the indicated proteins at concentrations ranging from 50 nM to 800 nM and incubated on ice for at least 10 min. Immediately before loading to a non-denaturing 5 % acrylamide gel (59:1 ratio of acrylamide to bis-acrylamide), sucrose was added to a final concentration of 4% (w/v). 0.2x TBE (17.5 mM Tris, pH 8.0, 17.5 mM boric acid, and 0.4 mM EDTA) was used as a running buffer. After electrophoresis, the DNA in the gel was stained by incubating it for 10 min at RT with SYBR™ Gold (Thermo Fisher) before imaging with a ChemiDoc imager (Bio-Rad). To evaluate affinity, the shift of either free nucleosomes or free DNA was observed.

### Protein Biotinylation

RB or the indicated constructs thereof were biotinylated in 50 mM KPO_4_, pH 7.7, 100 mM NaCl by the addition of a 27.5 x molar excess of EZ-Link™ NHS-PEG4-Biotin (Life Technologies). The reaction was incubated for 20 min at 4 °C and quenched by adding 100 mM ammonium bicarbonate. The reaction mixture was dialyzed against 3x 400 ml of 50 mM KPO_4_, pH 7.7, 100 mM NaCl, 1 mM DTT, 2 % glycerol for at least 2 h each. Biotinylated proteins were concentrated and used for further experiments, or flash-frozen in liquid nitrogen and stored at −80 °C.

### Biolayer Interferometry (BLI)

Biolayer interferometry (BLI) experiments were conducted using an Octet^®^ N1 System (Sartorius) at 20 °C. Octet® High Precision Streptavidin (SAX) Biosensors from Sartorius were hydrated for 10 min in reaction buffer (50 mM KPO_4_, pH 7.7, 100 mM NaCl) and then loaded with biotinylated RB constructs in the same buffer for 5 min. Free binding sites on the chip were blocked using 1 mM PEG_24_-biotin acid (Sigma-Aldrich) in reaction buffer.

The binding experiments with KDM5s were done in the reaction buffer. The baseline was measured for 10 seconds against the buffer before each measurement. Protein concentrations ranged from 1 nM to 4 µM, with association and dissociation times of 90 sec. To account for non-specific binding of KDM5s, each concentration was measured against a PEG_24_-biotin blocked chip and subtracted from the signal from the binding experiments.

K_D_ values were determined by plotting the local fit maximum response (nm) as a function of RB concentrations (nM) using Octet^®^ N1 Software 1.4.0.13. Global fitting was applied to all data, using a heterogeneous binding model, and the titration curves were fit using the following steady-state analysis equation: Response = R_max_ × conc/(K_D_+ conc), where R_max_ is the local fit maximum response, conc is the concentration of the RB, and K_D_ is the equilibrium dissociation constant. All experiments were repeated at least three times, and the average binding constants, along with their standard errors of the mean, were calculated in GraphPad Prism. Significance between conditions was calculated in GraphPad Prism using one-way ANOVA, with P<0.05 considered significant.

### Chemical cross-linking

2 µM of KDM5s in reaction buffer (50 mM KPO_4_, pH 7.7, 100 mM NaCl) was mixed with RB at a concentration range from 2 µM to 4 µM and incubated for at least 10 min on ice. BS^3^ or DSBU was added to a final concentration of 0.5 mM, and the reaction was mixed and incubated on ice for 20 min. The reaction was quenched by adding ammonium bicarbonate to a final concentration of 100 mM. Cross-linking was assessed by SDS-PAGE. Gel bands were cut out for in-gel digestion, and the remaining cross-linking solution was flash frozen and stored at - 80 °C until in-solution digestion.

### In-gel digestion (IGD) of cross-linked proteins

Gel regions containing cross-linked proteins were excised and washed twice each with water and 100 mM ammonium bicarbonate. The proteins were reduced with 10 mM TCEP (30 min, 62 °C) while gently shaking, and subsequently alkylated with 55 mM iodoacetamide (IAA) (30 min, room temperature, in the dark). The gel slabs were subsequently washed three times with 50 µl acetonitrile. After the last wash, the gel slabs were completely dried by using a vacuum concentrator (Eppendorf) for 5 min. The dried gel pieces were next incubated with 200 µl 10 ng/µl trypsin (37 °C, 16 h, vigorous shaking). The digestion was stopped by addition of formic acid to a final concentration of 5 % (v/v) formic acid. The digestion supernatant was transferred to a fresh Eppendorf tube. The remaining gel pieces were subsequently washed 3× with 50 µl acetonitrile. The supernatants of these washes were combined with the recovered digestion mix and dried in a vacuum concentrator (Eppendorf).

The cleared tryptic digests were then desalted on home-made C18 StageTips as described.^[75]^ Briefly, acidified peptides were immobilized and washed on a 2 disc C18 StageTip. After elution from the StageTips, samples were dried using a vacuum concentrator (Eppendorf) and the peptides were taken up in 0.1 % formic acid solution (10 μl) and directly used for LC-MS/MS experiments (see below for details).

### Single-pot, solid-phase-enhanced sample-preparation (SP3) of cross-linked proteins

The preparation of cross-linked samples for LC/MS/MS is based on the SP3 protocol.^[76]^ 30 µg of protein from each cross-linking sample was taken up in 100 µl 1× SP3 lysis buffer (final concentrations: 5 % (w/v) SDS; 10 mM TCEP; 200 μl 40 mM chloracetamide; 200 mM HEPES pH 8) and heated for 5 min at 90 °C. After cooling the samples to room temperature (on ice) a mix of 150 µg hydrophobic (#65152105050250) and 150 µg hydrophilic (#45152105050250) SeraMag Speed Beads (Cytiva) was added (bead to protein ratio 10 to 1) and gently mixed. Then 100 µl 100% v/v Ethanol (EtOH) was added before incubation for 20 min at 24 °C, shaking vigorously. The beads were collected on a magnet, and the supernatant was aspirated. The beads were then washed 4 times with 180 µl 80 % EtOH (collection time on the magnet minimum of 4 min). The beads were then finally taken up in 100 µl 25 mM ammonium bicarbonate (ABC) containing 1 µg Trypsin (Protein:Trypsin ratio 30:1). To help bead dissociation, samples were incubated for 1 min in a sonification bath (preheated to 37 °C). Samples were incubated overnight, shaking vigorously (1300 rpm). Samples were acidified with formic acid (FA, final 1 % (v/v)) before collection on a magnet. The supernatants were transferred to a fresh Eppendorf tube, before removing trace beads using a magnet for 5 min. The tryptic digests were then desalted on home-made C18 StageTips as described.^[75]^. After elution (80 % ACN; 0.1 % FA) samples were dried using a vacuum concentrator (Eppendorf) and the peptides were taken up in 0.1 % formic acid

### LC-MS/MS settings

MS Experiments were performed on an Orbitrap LUMOS instrument (Thermo) coupled to Vanquish Neo ultra-performance liquid chromatography (UPLC) system (Thermo). The UPLC was operated in the one-column mode. The analytical column was a fused silica capillary (75 µm × 28 cm) with an integrated fritted emitter (CoAnn Technologies) packed in-house with Kinetex 1.7 µm core shell beads (Phenomenex). The analytical column was encased by a column oven (Sonation PRSO-V2) and attached to a nanospray flex ion source (Thermo). The column oven temperature was set to 50 °C during sample loading and data acquisition. The LC was equipped with two mobile phases: solvent A (0.2 % FA, 2 % Acetonitrile, ACN, 97.8 % H_2_O) and solvent B (0.2 % FA, 80 % ACN, 19.8 % H_2_O). All solvents were of UPLC grade (Honeywell). Peptides were directly loaded onto the analytical column with a maximum flow rate that would not exceed the set pressure limit of 950 bar (usually around 0.4–0.6 µl/min). Peptides were subsequently separated on the analytical column by running a 67 min gradient of solvent A and solvent B (start with 8 % B; gradient 8 % to 40 % B for 50:00 min; gradient 40 % to 100 % B for 11:00 min; end with B 100 % for 6 min) at a flow rate of 250 nl/min. The mass spectrometer was controlled by the Orbitrap Fusion Lumos Tune Application (version 4.1.4244) and operated using the Xcalibur software (version v4.7.69.37). The mass spectrometer was set in the positive ion mode. The ionization potential (spray voltage) was set to 2.3 kV. Source fragmentation was turned off. Precursor ion scanning was performed in the Orbitrap analyzer (FT; fourier transform mass spectrometer) in the scan range of m/z 380-1400 and at a resolution of 60000 with the internal lock mass option turned on (lock mass was 445.120025 m/z, polysiloxane).^[77]^ AGC (automatic gain control) was set to “standard” and acquisition time to “auto”. Product ion spectra were recorded in a data dependent fashion in the Orbitrap at a variable scan range (“auto”) and at a resolution of 30000. Peptides were analyzed using a “top speed” regime (repeating cycle of full precursor ion scan (AGC target 200 %; 100000 ions; acquisition time 70 ms) followed by further dependent MS2 scans for 3 s (minimum intensity threshold 5×10^4^)). The MS2 precursor ions were isolated using the quadrupole (isolation window 1.6 m/z), and fragmentation was achieved by Higher-energy C-trap dissociation (HCD) (normalized collision mode set to “assisted” and normalized collision energy set to 25, 30, 40). During MS2 data acquisition dynamic ion exclusion was set to 30 s. Only charge states between 3-8 were considered for fragmentation. Detailed LC-MS settings can be found in the supplementary file Sample_Legend_and_LC-MS_Settings_v01.docx.

### Data Processing Protocol CL-MS

The searches for cross-linked peptides were performed with MetaMorpheus (MM) (version 1.0.6.).^[78]^ Searches were performed on the Thermo RAW files using the database ACE_0945_0950_SOI_plus_con_v01.fasta. The database contains the sequences for the proteins of interest and known contaminating proteins often found in MS samples. The peptide spectrum match FDR for MM was 0.01 (based on target-decoy approach, decoys are generated by the software). The settings for MetaMorpheus were: cross-linker name = DSS (note: DSS and BS^3^ have identical cross-linker size; cross-linker type = non-cleavable; cross-linker mass = 138.06808; cross-linker modification site 1 = K; cross-linker modification site 2 = K) or DSBU (cross-linker type = cleavable; cross-linker mass = 196.0848; cross-linker modification site 1 = K; cross-linker modification site 2 = K or KSTY); protease = trypsin; maximum missed cleavages = 3; minimum peptide length = 5; maximum peptide length = 60; initiator methionine behavior = Variable; max modification isoforms = 1024; fixed modifications = Carbamidomethyl on C, variable modifications = Oxidation on M; parent mass tolerance(s) = ±10 ppm; product mass tolerance = ±20 ppm. The results from the searches were converted to the ProXL XML format using the converter metaMorph2ProxlXML.jar (from the ProXL website; https://proxl-ms.org/) and uploaded to a local installation of the ProXL server.^[79]^ Analysis and evaluation of cross-links was performed on our local ProXL Server.

### Negative-stain sample preparation and data collection

400 mesh copper grids with continuous carbon (EM Sciences) were glow-discharged with a Tergeo Plasma Cleaner (PIE Scientific). 4 µl of 25-50 nM KDM5 in 50 mM KPO_4_, pH 7.7, 100 mM NaCl were incubated for 30 s on the grids. To avoid formation of uranyl phosphate crystals, the excess sample was side blotted, and the grids were incubated twice with 50 mM HEPES, pH 7.7, 100 mM NaCl for 1 sec each. Samples were stained by adding 4×2 % uranyl formate solution for increasing incubation times (2, 4, 8, 16 s). After a final 10-second incubation, excess stain was removed by blotting with filter paper.

Grids were imaged using a Talos L120C microscope (Thermo Fisher) operated at 120 keV equipped with a Ceta 16M detector (Thermo Fisher). Micrographs were acquired at a nominal pixel size of 1.52 Å and total dose of 25-30 e^−^ per Å^2^.

### EM data processing and docking of atomic models

All processing was done using CryoSPARC.^[80]^ Contrast transfer function (CTF) estimates were performed with CTFFIND4^[81]^, blob picker was used for particle picking and iterative rounds of reference-free 2D classification. Classes representing damaged protein or artifacts were removed in several rounds of 2D classification. An *ab initio* approach was performed to generate a template for template picking. Low-quality or false particle picks were filtered out using 2D classification. *Ab initio* 3D reconstructions were generated, followed by homogeneous refinement (Suppl. Fig. 3,4).

AlphaFold3 predictions for KDM5A and KDM5B (Fig. 1 B,C) were manually rigid-body fitted into the obtained negative-staining volumes using ChimeraX.^[82,83]^ To account for flexibility, AlphaFold3 predictions were separated into two structures: for KDM5A, aa 1-795 and aa 796-1690; and for KDM5B, aa 1-755 and aa 756-1544. Alphafold3 statistics are shown in Suppl. Fig. 5.

### Cross-linking validation

For KDM5A, Cɑ-distances between cross-linked aa were highlighted in the crystal structure from Vinogradova *et al.*^[52]^ (PDB 5CEH), and for RB, they are highlighted in the structure of Burke *et al.*^[53]^ (PDB: 4ELJ). Cross-links outside of the sequence from the crystal structure are not shown. Distances were calculated using ChimeraX. Based on the linker length of BS^3^ (11.4 Å) and DSBU (12.5 Å), a Cɑ-distance of less than 24 Å was deemed reasonable (highlighted in green), 24-30 Å as possible (yellow), and a Cɑ-distance above 30 Å as unlikely (red).^[84]^

### Generation of Alphafold predictions

Atomic models of KDM5A and KDM5B were generated using the Alphafold Server from Google LLC.^[85]^ Sequences used for predictions were obtained from Uniprot (KDM5A: P29375, KDM5B: Q9UGL1). Prediction statistics and atomic models were prepared using ChimeraX.^[82,83]^ (Suppl. Fig. 5) PAE plots were generated using PAE Viewer.^[86]^

### Sequence alignment and disorder predictions

The sequence alignment of KDM5A and KDM5B was generated using CLUSTALW.^[87]^ Disorder predictions were done using PONDR.^[88–90]^ Sequences used for predictions were obtained from Uniprot (KDM5A: P29375, KDM5B: Q9UGL1). Alignment statistics and illustrations were analyzed with Jalview.^[91]^ (Suppl. Fig. 1B) Plots were generated using GraphPad Prism.

## References

[1] A. R. Mansisidor, V. I. Risca, “Chromatin accessibility: methods, mechanisms, and biological insights” Nucleus 2022, 13, 238–278.

[2] T. Kouzarides, “Chromatin Modifications and Their Function” Cell 2007, 128, 693– 705.

[3] M. Yun, J. Wu, J. L. Workman, B. Li, “Readers of histone modifications” Cell Res. 2011, 21, 564–578.

[4] H. Yu, B. J. Lesch, “Functional Roles of H3K4 Methylation in Transcriptional Regulation” *Mol. Cell*. Biol. 2024, 44, 505–515.

[5] R. J. Klose, Q. Yan, Z. Tothova, K. Yamane, H. Erdjument-Bromage, P. Tempst, D. G. Gilliland, Y. Zhang, W. G. Kaelin, “The Retinoblastoma Binding Protein RBP2 Is an H3K4 Demethylase” Cell 2007, 128, 889–900.

[6] Y. Xiang, Z. Zhu, G. Han, X. Ye, B. Xu, Z. Peng, Y. Ma, Y. Yu, H. Lin, A. P. Chen, C. D. Chen, “JARID1B is a histone H3 lysine 4 demethylase up-regulated in prostate cancer” *Proc*. Natl. Acad. Sci. 2007, 104, 19226–19231.

[7] M. Tahiliani, P. Mei, R. Fang, T. Leonor, M. Rutenberg, F. Shimizu, J. Li, A. Rao, Y. Shi, “The histone H3K4 demethylase SMCX links REST target genes to X-linked mental retardation” Nature 2007, 447, 601–605.

[8] E. Pavlenko, T. Ruengeler, P. Engel, S. Poepsel, “Functions and Interactions of Mammalian KDM5 Demethylases” *Front*. Genet. 2022, 13, 906662.

[9] D. J. Seward, G. Cubberley, S. Kim, M. Schonewald, L. Zhang, B. Tripet, D. L. Bentley, “Demethylation of trimethylated histone H3 Lys4 in vivo by JARID1 JmjC proteins” *Nat. Struct*. Mol. Biol. 2007, 14, 240–242.

[10] S. Jamshidi, S. Catchpole, J. Chen, C. So, J. Burchell, K. Rahman, J. Taylor-papadimitriou, “KDM5B protein expressed in viable and fertile ΔARID mice exhibit no demethylase activity” *Int*. J. Oncol. 2021, 59, 96.

[11] F. S. Ugur, M. J. S. Kelly, D. G. Fujimori, “Chromatin Sensing by the Auxiliary Domains of KDM5C Regulates Its Demethylase Activity and Is Disrupted by X-linked Intellectual Disability Mutations” *J*. Mol. Biol. 2023, 435, 167913.

[12] A. Kataria, S. Tyagi, “Domain architecture and protein–protein interactions regulate KDM5A recruitment to the chromatin” Epigenetics 2023, 18, 2268813.

[13] J. R. Horton, A. Engstrom, E. L. Zoeller, X. Liu, J. R. Shanks, X. Zhang, M. A. Johns, P. M. Vertino, H. Fu, X. Cheng, “Characterization of a Linked Jumonji Domain of the KDM5/JARID1 Family of Histone H3 Lysine 4 Demethylases” *J*. Biol. Chem. 2016, 291, 2631–2646.

[14] I. O. Torres, K. M. Kuchenbecker, C. I. Nnadi, R. J. Fletterick, M. J. S. Kelly, D. G. Fujimori, “Histone demethylase KDM5A is regulated by its reader domain through a positive-feedback mechanism” *Nat*. Commun. 2015, 6, 6204.

[15] J. E. Longbotham, C. M. Chio, V. Dharmarajan, M. J. Trnka, I. O. Torres, D. Goswami, K. Ruiz, A. L. Burlingame, P. R. Griffin, D. G. Fujimori, “Histone H3 binding to the PHD1 domain of histone demethylase KDM5A enables active site remodeling” *Nat*. Commun. 2019, 10, 94.

[16] B. J. Klein, L. Piao, Y. Xi, H. Rincon-Arano, S. B. Rothbart, D. Peng, H. Wen, C. Larson, X. Zhang, X. Zheng, M. A. Cortazar, P. V. Peña, A. Mangan, D. L. Bentley, B. D. Strahl, M. Groudine, W. Li, X. Shi, T. G. Kutateladze, “The Histone-H3K4-Specific Demethylase KDM5B Binds to Its Substrate and Product through Distinct PHD Fingers” Cell Rep. 2014, 6, 325–335.

[17] G. G. Wang, J. Song, Z. Wang, H. L. Dormann, F. Casadio, H. Li, J.-L. Luo, D. J. Patel, C. D. Allis, “Haematopoietic malignancies caused by dysregulation of a chromatin-binding PHD finger” Nature 2009, 459, 847–851.

[18] A. G. Scibetta, S. Santangelo, J. Coleman, D. Hall, T. Chaplin, J. Copier, S. Catchpole, J. Burchell, J. Taylor-Papadimitriou, “Functional Analysis of the Transcription Repressor PLU-1/JARID1B” *Mol. Cell*. Biol. 2007, 27, 7220–7235.

[19] S. Tu, Y.-C. Teng, C. Yuan, Y.-T. Wu, M.-Y. Chan, A.-N. Cheng, P.-H. Lin, L.-J. Juan, M.-D. Tsai, “The ARID domain of the H3K4 demethylase RBP2 binds to a DNA CCGCCC motif” *Nat. Struct*. Mol. Biol. 2008, 15, 419–421.

[20] A.-S. B. Brier, A. Loft, J. G. S. Madsen, T. Rosengren, R. Nielsen, S. F. Schmidt, Z. Liu, Q. Yan, H. Gronemeyer, S. Mandrup, “The KDM5 family is required for activation of pro-proliferative cell cycle genes during adipocyte differentiation” Nucleic Acids Res. 2017, 45, 1743–1759.

[21] M. Uhlén, L. Fagerberg, B. M. Hallström, C. Lindskog, P. Oksvold, A. Mardinoglu, Å. Sivertsson, C. Kampf, E. Sjöstedt, A. Asplund, I. Olsson, K. Edlund, E. Lundberg, S. Navani, C. A.-K. Szigyarto, J. Odeberg, D. Djureinovic, J. O. Takanen, S. Hober, T. Alm, P.-H. Edqvist, H. Berling, H. Tegel, J. Mulder, J. Rockberg, P. Nilsson, J. M. Schwenk, M. Hamsten, K. Von Feilitzen, M. Forsberg, L. Persson, F. Johansson, M. Zwahlen, G. Von Heijne, J. Nielsen, F. Pontén, “Tissue-based map of the human proteome” Science 2015, 347, 1260419.

[22] A. Barrett, B. Madsen, J. Copier, P. J. Lu, L. Cooper, A. G. Scibetta, J. Burchell, J. Taylor-Papadimitriou, “PLU-1 nuclear protein, which is upregulated in breast cancer, shows restricted expression in normal human adult tissues: A new cancer/testis antigen?” Int. J. Cancer 2002, 101, 581–588.

[23] J. Yoo, G. W. Kim, Y. H. Jeon, J. Y. Kim, S. W. Lee, S. H. Kwon, “Drawing a line between histone demethylase KDM5A and KDM5B: their roles in development and tumorigenesis” *Exp*. Mol. Med. 2022, 54, 2107–2117.

[24] Burchell, “PLU-1/JARID1B/KDM5B is required for embryonic survival and contributes to cell proliferation in the mammary gland and in ER+ breast cancer cells” *Int*. J. Oncol. 2011, 38, DOI 10.3892/ijo.2011.956.

[25] L. El Hayek, I. O. Tuncay, N. Nijem, J. Russell, S. Ludwig, K. Kaur, X. Li, P. Anderton, M. Tang, A. Gerard, A. Heinze, P. Zacher, H. S. Alsaif, A. Rad, K. Hassanpour, M. R. Abbaszadegan, C. Washington, B. R. DuPont, R. J. Louie, CAUSES Study, M. Couse, M. Faden, R. C. Rogers, R. Abou Jamra, E. R. Elias, R. Maroofian, H. Houlden, A. Lehman, B. Beutler, M. H. Chahrour, “KDM5A mutations identified in autism spectrum disorder using forward genetics” eLife 2020, 9, e56883.

[26] S. Kong, W. Kim, H. Lee, H. Kim, “The histone demethylase KDM5A is required for the repression of astrocytogenesis and regulated by the translational machinery in neural progenitor cells” FASEB J. 2018, 32, 1108–1119.

[27] S. U. Schmitz, M. Albert, M. Malatesta, L. Morey, J. V. Johansen, M. Bak, N. Tommerup, I. Abarrategui, K. Helin, “Jarid1b targets genes regulating development and is involved in neural differentiation” EMBO J. 2011, 30, 4586–4600.

[28] A. Rojas, R. Aguilar, B. Henriquez, J. B. Lian, J. L. Stein, G. S. Stein, A. J. Van Wijnen, B. Van Zundert, M. L. Allende, M. Montecino, “Epigenetic Control of the Bone-master Runx2 Gene during Osteoblast-lineage Commitment by the Histone Demethylase JARID1B/KDM5B” *J*. Biol. Chem. 2015, 290, 28329–28342.

[29] R. Váraljai, A. B. M. M. K. Islam, M. L. Beshiri, J. Rehman, N. Lopez-Bigas, E. V. Benevolenskaya, “Increased mitochondrial function downstream from KDM5A histone demethylase rescues differentiation in pRB-deficient cells” Genes Dev. 2015, 29, 1817–1834.

[30] D. Defeo-Jones, P. S. Huang, R. E. Jones, K. M. Haskell, G. A. Vuocolo, M. G. Hanobik, H. E. Huber, A. Oliff, “Cloning of cDNAs for cellular proteins that bind to the retinoblastoma gene product” Nature 1991, 352, 251–254.

[31] R. A. Weinberg, “The retinoblastoma protein and cell cycle control” Cell 1995, 81, 323–330.

[32] S. P. Chellappan, S. Hiebert, M. Mudryj, J. M. Horowitz, J. R. Nevins, “The E2F transcription factor is a cellular target for the RB protein” Cell 1991, 65, 1053–1061.

[33] P.-L. Chen, P. Scully, J.-Y. Shew, J. Y. J. Wang, W.-H. Lee, “Phosphorylation of the retinoblastoma gene product is modulated during the cell cycle and cellular differentiation” Cell 1989, 58, 1193–1198.

[34] L. Zhou, D. S.-C. Ng, J. C. Yam, L. J. Chen, C. C. Tham, C. P. Pang, W. K. Chu, “Post-translational modifications on the retinoblastoma protein” *J*. Biomed. Sci. 2022, 29, 33.

[35] M. Gao, H. Li, J. Zhang, “RB functions as a key regulator of senescence and tumor suppression” *Semin*. Cancer Biol. 2025, 109, 1–7.

[36] F. A. Dick, D. W. Goodrich, J. Sage, N. J. Dyson, “Non-canonical functions of the RB protein in cancer” *Nat. Rev*. Cancer 2018, 18, 442–451.

[37] A. Chicas, A. Kapoor, X. Wang, O. Aksoy, A. G. Evertts, M. Q. Zhang, B. A. Garcia, E. Bernstein, S. W. Lowe, “H3K4 demethylation by Jarid1a and Jarid1b contributes to retinoblastoma-mediated gene silencing during cellular senescence” *Proc*. Natl. Acad. Sci. 2012, 109, 8971–8976.

[38] J. H. Nijwening, E.-J. Geutjes, R. Bernards, R. L. Beijersbergen, “The Histone Demethylase Jarid1b (Kdm5b) Is a Novel Component of the Rb Pathway and Associates with E2f-Target Genes in MEFs during Senescence” PLoS ONE 2011, 6, e25235.

[39] M. L. Beshiri, K. B. Holmes, W. F. Richter, S. Hess, A. B. M. M. K. Islam, Q. Yan, L. Plante, L. Litovchick, N. Gévry, N. Lopez-Bigas, W. G. Kaelin, E. V. Benevolenskaya, “Coordinated repression of cell cycle genes by KDM5A and E2F4 during differentiation” *Proc*. Natl. Acad. Sci. 2012, 109, 18499–18504.

[40] E. V. Benevolenskaya, H. L. Murray, P. Branton, R. A. Young, W. G. Kaelin, “Binding of pRB to the PHD Protein RBP2 Promotes Cellular Differentiation” *Mol*. Cell 2005, 18, 623–635.

[41] W. Lin, J. Cao, J. Liu, M. L. Beshiri, Y. Fujiwara, J. Francis, A. D. Cherniack, C. Geisen, L. P. Blair, M. R. Zou, X. Shen, D. Kawamori, Z. Liu, C. Grisanzio, H. Watanabe, Y. A. Minamishima, Q. Zhang, R. N. Kulkarni, S. Signoretti, S. J. Rodig, R. T. Bronson, S. H. Orkin, D. P. Tuck, E. V. Benevolenskaya, M. Meyerson, W. G. Kaelin, Q. Yan, “Loss of the retinoblastoma binding protein 2 (RBP2) histone demethylase suppresses tumorigenesis in mice lacking *Rb1* or *Men1*” *Proc*. Natl. Acad. Sci. 2011, 108, 13379–13386.

[42] A. Roesch, B. Becker, W. Schneider-Brachert, I. Hagen, M. Landthaler, T. Vogt, “Re-Expression of the Retinoblastoma-Binding Protein 2-Homolog 1 Reveals Tumor-Suppressive Functions in Highly Metastatic Melanoma Cells” *J*. Invest. Dermatol. 2006, 126, 1850–1859.

[43] A. Roesch, M. Fukunaga-Kalabis, E. C. Schmidt, S. E. Zabierowski, P. A. Brafford, A. Vultur, D. Basu, P. Gimotty, T. Vogt, M. Herlyn, “A Temporarily Distinct Subpopulation of Slow-Cycling Melanoma Cells Is Required for Continuous Tumor Growth” SCell 2010, 141, 583–594.

[44] J.-O. Lee, A. A. Russo, N. P. Pavletich, “Structure of the retinoblastoma tumour-suppressor pocket domain bound to a peptide from HPV E7” Nature 1998, 391, 859–865.

[45] M. G. Andrusiak, R. Vandenbosch, F. A. Dick, D. S. Park, R. S. Slack, “LXCXE- independent chromatin remodeling by Rb/E2f mediates neuronal quiescence” Cell Cycle 2013, 12, 1416–1423.

[46] Y. W. Kim, G. A. Otterson, R. A. Kratzke, A. B. Coxon, F. J. Kaye, “Differential Specificity for Binding of Retinoblastoma Binding Protein 2 to RB, pl07, and TATA-Binding Protein” *Mol. Cell*. Biol. 1994, 14, 7256–7264.

[47] Z. U. Zargar, M. R. Kimidi, S. Tyagi, “Dynamic site-specific recruitment of RBP2 by pocket protein p130 modulates H3K4 methylation on E2F-responsive promoters” Nucleic Acids Res. 2018, 46, 174–188.

[48] A. Roesch, B. Becker, S. Meyer, P. Wild, C. Hafner, M. Landthaler, T. Vogt, “Retinoblastoma-binding protein 2-homolog 1: a retinoblastoma-binding protein downregulated in malignant melanomas” *Mod*. Pathol. 2005, 18, 1249–1257.

[49] A. M. Palla, C.-C. Lin, M. J. Trnka, E. M. Leao, N. Petronikolou, A. L. Burlingame, R. K. McGinty, D. G. Fujimori, “An Intrinsically Disordered Region of Histone Demethylase KDM5A Activates Catalysis Through Interactions With the Nucleosomal Acidic Patch and DNA” *J*. Mol. Biol. 2025, 437, 169301.

[50] J. Dorosz, L. H. Kristensen, N. G. Aduri, O. Mirza, R. Lousen, S. Bucciarelli, V. Mehta, S. Sellés-Baiget, S. M. Ø. Solbak, A. Bach, P. Mesa, P. A. Hernandez, G. Montoya, T. T. T. N. Nguyen, K. D. Rand, T. Boesen, M. Gajhede, “Molecular architecture of the Jumonji C family histone demethylase KDM5B” *Sci*. Rep. 2019, 9, 4019.

[51] M. D. Simon, F. Chu, L. R. Racki, C. C. De La Cruz, A. L. Burlingame, B. Panning, G. J. Narlikar, K. M. Shokat, “The Site-Specific Installation of Methyl-Lysine Analogs into Recombinant Histones” Cell 2007, 128, 1003–1012.

[52] M. Vinogradova, V. S. Gehling, A. Gustafson, S. Arora, C. A. Tindell, C. Wilson, K. E. Williamson, G. D. Guler, P. Gangurde, W. Manieri, J. Busby, E. M. Flynn, F. Lan, H. Kim, S. Odate, A. G. Cochran, Y. Liu, M. Wongchenko, Y. Yang, T. K. Cheung, T. M. Maile, T. Lau, M. Costa, G. V. Hegde, E. Jackson, R. Pitti, D. Arnott, C. Bailey, S. Bellon, R. T. Cummings, B. K. Albrecht, J.-C. Harmange, J. R. Kiefer, P. Trojer, M. Classon, “An inhibitor of KDM5 demethylases reduces survival of drug-tolerant cancer cells” *Nat*. Chem. Biol. 2016, 12, 531–538.

[53] J. R. Burke, G. L. Hura, S. M. Rubin “Structures of inactive retinoblastoma protein reveal multiple mechanisms for cell cycle control” Genes Dev. 2012, 26, 1156–1166.

[54] S. Putta, L. Alvarez, S. Lüdtke, P. Sehr, G. A. Müller, S. M. Fernandez, S. Tripathi, J. Lewis, T. J. Gibson, L. B. Chemes, S. M. Rubin, “Structural basis for tunable affinity and specificity of LxCxE-dependent protein interactions with the retinoblastoma protein family” Structure 2022, 30, 1340–1353.e3.

[55] J. R. Horton, X. Liu, M. Gale, L. Wu, J. R. Shanks, X. Zhang, P. J. Webber, J. S. K. Bell, S. C. Kales, B. T. Mott, G. Rai, D. J. Jansen, M. J. Henderson, D. J. Urban, M. D. Hall, A. Simeonov, D. J. Maloney, M. A. Johns, H. Fu, A. Jadhav, P. M. Vertino, Q. Yan, X. Cheng, “Structural Basis for KDM5A Histone Lysine Demethylase Inhibition by Diverse Compounds” Cell Chem. Biol. 2016, 23, 769–781.

[56] C. J. Spangler, A. Skrajna, C. A. Foley, A. Nguyen, G. R. Budziszewski, D. N. Azzam, E. C. Arteaga, H. C. Simmons, C. B. Smith, N. A. Wesley, E. M. Wilkerson, J.-M. E. McPherson, D. Kireev, L. I. James, S. V. Frye, D. Goldfarb, R. K. McGinty, “Structural basis of paralog-specific KDM2A/B nucleosome recognition” *Nat*. Chem. Biol. 2023, 19, 624–632.

[57] P. Zhang, K. Torres, X. Liu, C. Liu, R. E. Pollock, “An Overview of Chromatin-Regulating Proteins in Cells” *Curr*. Protein Pept. Sci. 2016, 17, 401–410.

[58] T. Vogt, M. Kroiss, M. McClelland, C. Gruss, B. Becker, A. K. Bosserhoff, G. Rumpler, T. Bogenrieder, M. Landthaler, W. Stolz, “Deficiency of a novel retinoblastoma binding protein 2-homolog is a consistent feature of sporadic human melanoma skin cancer” *Lab*. Investig. J. Tech. Methods Pathol. 1999, 79, 1615–1627.

[59] A. Roesch, A. M. Mueller, T. Stempfl, C. Moehle, M. Landthaler, T. Vogt, “RBP2- H1/JARID1B is a transcriptional regulator with a tumor suppressive potential in melanoma cells” *Int*. J. Cancer 2008, 122, 1047–1057.

[60] Q. Li, L. Shi, B. Gui, W. Yu, J. Wang, D. Zhang, X. Han, Z. Yao, Y. Shang, “Binding of the JmjC Demethylase JARID1B to LSD1/NuRD Suppresses Angiogenesis and Metastasis in Breast Cancer Cells by Repressing Chemokine CCL14” Cancer Res. 2011, 71, 6899–6908.

[61] A. Brehm, S. J. Nielsen, E. A. Miska, D. J. McCance, J. L. Reid, A. J. Bannister, T. Kouzarides, “The E7 oncoprotein associates with Mi2 and histone deacetylase activity to promote cell growth” EMBO J. 1999, 18, 2449–2458.

[62] J. B. Rayman, Y. Takahashi, V. B. Indjeian, J.-H. Dannenberg, S. Catchpole, R. J. Watson, H. Te Riele, B. D. Dynlacht, “E2F mediates cell cycle-dependent transcriptional repression in vivo by recruitment of an HDAC1/mSin3B corepressor complex” Genes Dev. 2002, 16, 933–947.

[63] N. Lopez-Bigas, T. A. Kisiel, D. C. DeWaal, K. B. Holmes, T. L. Volkert, S. Gupta, J. Love, H. L. Murray, R. A. Young, E. V. Benevolenskaya, “Genome-wide Analysis of the H3K4 Histone Demethylase RBP2 Reveals a Transcriptional Program Controlling Differentiation” *Mol*. Cell 2008, 31, 520–530.

[64] E. S. Knudsen, J. Y. J. Wang, “Differential Regulation of Retinoblastoma Protein Function by Specific Cdk Phosphorylation Sites” *J*. Biol. Chem. 1996, 271, 8313–8320.

[65] The wwPDB Consortium, J. Turner, S. Abbott, N. Fonseca, R. Pye, L. Carrijo, A. K. Duraisamy, O. Salih, Z. Wang, G. J. Kleywegt, K. L. Morris, A. Patwardhan, S. K. Burley, G. Crichlow, Z. Feng, J. W. Flatt, S. Ghosh, B. P. Hudson, C. L. Lawson, Y. Liang, E. Peisach, I. Persikova, M. Sekharan, C. Shao, J. Young, S. Velankar, D. Armstrong, M. Bage, W. M. Bueno, G. Evans, R. Gaborova, S. Ganguly, D. Gupta, D. Harrus, A. Tanweer, M. Bansal, V. Rangannan, G. Kurisu, H. Cho, Y. Ikegawa, Y. Kengaku, J. Y. Kim, S. Niwa, J. Sato, A. Takuwa, J. Yu, J. C. Hoch, K. Baskaran, W. Xu, W. Zhang, X. Ma, “EMDB—the Electron Microscopy Data Bank” Nucleic Acids Res. 2024, 52, D456–D465.

[66] J. A. Vizcaíno, A. Csordas, N. del-Toro, J. A. Dianes, J. Griss, I. Lavidas, G. Mayer, Y. Perez-Riverol, F. Reisinger, T. Ternent, Q.-W. Xu, R. Wang, H. Hermjakob, “2016 update of the PRIDE database and its related tools” Nucleic Acids Res. 2016, 44, 11033–11033.

[67] P. N. Dyer, R. S. Edayathumangalam, C. L. White, Y. Bao, S. Chakravarthy, U. M. Muthurajan, K. Luger in Methods Enzymol., Elsevier, 2003, pp. 23–44.

[68] T. W. Roy, A. S. Bhagwat “Kinetic studies of Escherichia coli AlkB using a new fluorescence-based assay for DNA demethylation” Nucleic Acids Res. 2007, 35, e147–e147.

[69] J. Neuhold, K. Radakovics, A. Lehner, F. Weissmann, M. Q. Garcia, M. C. Romero, N. S. Berrow, P. Stolt-Bergner, “GoldenBac: a simple, highly efficient, and widely applicable system for construction of multi-gene expression vectors for use with the baculovirus expression vector system” BMC Biotechnol. 2020, 20, 26.

[70] P. Friedhoff, O. Gimadutdinow, T. Ruter, W. Wende, C. Urbanke, H. Thole, A. Pingoud, “A Procedure for Renaturation and Purification of the Extracellular Serratia marcescens Nuclease from Genetically Engineered Escherichia coli” Protein Expr. Purif. 1994, 5, 37–43.

[71] W. R. Sellers, B. G. Novitch, S. Miyake, A. Heith, G. A. Otterson, F. J. Kaye, A. B. Lassar, W. G. Kaelin, “Stable binding to E2F is not required for the retinoblastoma protein to activate transcription, promote differentiation, and suppress tumor cell growth” Genes Dev. 1998, 12, 95–106.

[72] K. Luger, T. J. Rechsteiner, A. J. Flaus, M. M. Y. Waye, T. J. Richmond, “Characterization of nucleosome core particles containing histone proteins made in bacteria 1 1Edited by A. Klug” J. Mol. Biol. 1997, 272, 301–311.

[73] P. T. Lowary, J. Widom, “New DNA sequence rules for high affinity binding to histone octamer and sequence-directed nucleosome positioning” *J*. Mol. Biol. 1998, 276, 19–42.

[74] S. Poepsel, V. Kasinath, E. Nogales, “Cryo-EM structures of PRC2 simultaneously engaged with two functionally distinct nucleosomes” *Nat. Struct*. Mol. Biol. 2018, 25, 154–162.

[75] J. Rappsilber, M. Mann, Y. Ishihama, “Protocol for micro-purification, enrichment, pre-fractionation and storage of peptides for proteomics using StageTips” *Nat*. Protoc. 2007, 2, 1896–1906.

[76] C. S. Hughes, S. Moggridge, T. Müller, P. H. Sorensen, G. B. Morin, J. Krijgsveld, “Single-pot, solid-phase-enhanced sample preparation for proteomics experiments” *Nat*. Protoc. 2019, 14, 68–85.

[77] J. V. Olsen, L. M. F. De Godoy, G. Li, B. Macek, P. Mortensen, R. Pesch, A. Makarov, O. Lange, S. Horning, M. Mann, “Parts per Million Mass Accuracy on an Orbitrap Mass Spectrometer via Lock Mass Injection into a C-trap” *Mol. Cell*. Proteomics 2005, 4, 2010–2021.

[78] L. Lu, R. J. Millikin, S. K. Solntsev, Z. Rolfs, M. Scalf, M. R. Shortreed, L. M. Smith, “Identification of MS-Cleavable and Noncleavable Chemically Cross-Linked Peptides with MetaMorpheus” J. Proteome Res. 2018, 17, 2370–2376.

[79] M. Riffle, D. Jaschob, A. Zelter, T. N. Davis, “ProXL (Protein Cross-Linking Database): A Platform for Analysis, Visualization, and Sharing of Protein Cross-Linking Mass Spectrometry Data” J. Proteome Res. 2016, 15, 2863–2870.

[80] A. Punjani, J. L. Rubinstein, D. J. Fleet, M. A. Brubaker, “cryoSPARC: algorithms for rapid unsupervised cryo-EM structure determination” Nat. Methods 2017, 14, 290–296.

[81] A. Rohou, N. Grigorieff, “CTFFIND4: Fast and accurate defocus estimation from electron micrographs” J. Struct. Biol. 2015, 192, 216–221.

[82] E. C. Meng, T. D. Goddard, E. F. Pettersen, G. S. Couch, Z. J. Pearson, J. H. Morris, T. E. Ferrin, “UCSF CHIMERAX: Tools for structure building and analysis” Protein Sci. 2023, 32, e4792.

[83] E. F. Pettersen, T. D. Goddard, C. C. Huang, E. C. Meng, G. S. Couch, T. I. Croll, J. H. Morris, T. E. Ferrin, “UCSF CHIMERAX: Structure visualization for researchers, educators, and developers” Protein Sci. 2021, 30, 70–82.

[84] E. D. Merkley, S. Rysavy, A. Kahraman, R. P. Hafen, V. Daggett, J. N. Adkins, “Distance restraints from crosslinking mass spectrometry: Mining a molecular dynamics simulation database to evaluate lysine–lysine distances” Protein Sci. 2014, 23, 747–759.

[85] J. Abramson, J. Adler, J. Dunger, R. Evans, T. Green, A. Pritzel, O. Ronneberger, L. Willmore, A. J. Ballard, J. Bambrick, S. W. Bodenstein, D. A. Evans, C.-C. Hung, M. O’Neill, D. Reiman, K. Tunyasuvunakool, Z. Wu, A. Žemgulytė, E. Arvaniti, C. Beattie, O. Bertolli, A. Bridgland, A. Cherepanov, M. Congreve, A. I. Cowen-Rivers, A. Cowie, M. Figurnov, F. B. Fuchs, H. Gladman, R. Jain, Y. A. Khan, C. M. R. Low, K. Perlin, A. Potapenko, P. Savy, S. Singh, A. Stecula, A. Thillaisundaram, C. Tong, S. Yakneen, E. D. Zhong, M. Zielinski, A. Žídek, V. Bapst, P. Kohli, M. Jaderberg, D. Hassabis, J. M. Jumper, “Accurate structure prediction of biomolecular interactions with AlphaFold 3” Nature 2024, 630, 493–500.

[86] C. Elfmann, J. Stülke, “PAE viewer: a webserver for the interactive visualization of the predicted aligned error for multimer structure predictions and crosslinks” Nucleic Acids Res. 2023, 51, W404–W410.

[87] J. D. Thompson, D. G. Higgins, T. J. Gibson “CLUSTAL W: improving the sensitivity of progressive multiple sequence alignment through sequence weighting, position-specific gap penalties and weight matrix choice” Nucleic Acids Res. 1994, 22, 4673–4680.

[88] K. Peng, S. Vucetic, P. Radivojac, C. J. Brown, A. K. Dunker, Z. Obradovic, “OPTIMIZING LONG INTRINSIC DISORDER PREDICTORS WITH PROTEIN EVOLUTIONARY INFORMATION” J. Bioinform. Comput. Biol. 2005, 03, 35–60.

[89] K. Peng, P. Radivojac, S. Vucetic, A. K. Dunker, Z. Obradovic, “Length-dependent prediction of protein intrinsic disorder” BMC Bioinformatics 2006, 7, 208.

[90] P. Romero, Z. Obradovic, X. Li, E. C. Garner, C. J. Brown, A. K. Dunker, “Sequence complexity of disordered protein” *Proteins Struct*. Funct. Genet. 2001, 42, 38–48.

[91] A. M. Waterhouse, J. B. Procter, D. M. A. Martin, M. Clamp, G. J. Barton, “Jalview Version 2—a multiple sequence alignment editor and analysis workbench” Bioinformatics 2009, 25, 1189–1191.

