## supplementary figures 1-9 for "Differences in substrate engagement and Retinoblastoma protein (RB) binding of human KDM5A and KDM5B"

### Supplementary Figure/Tables Titles and Legends

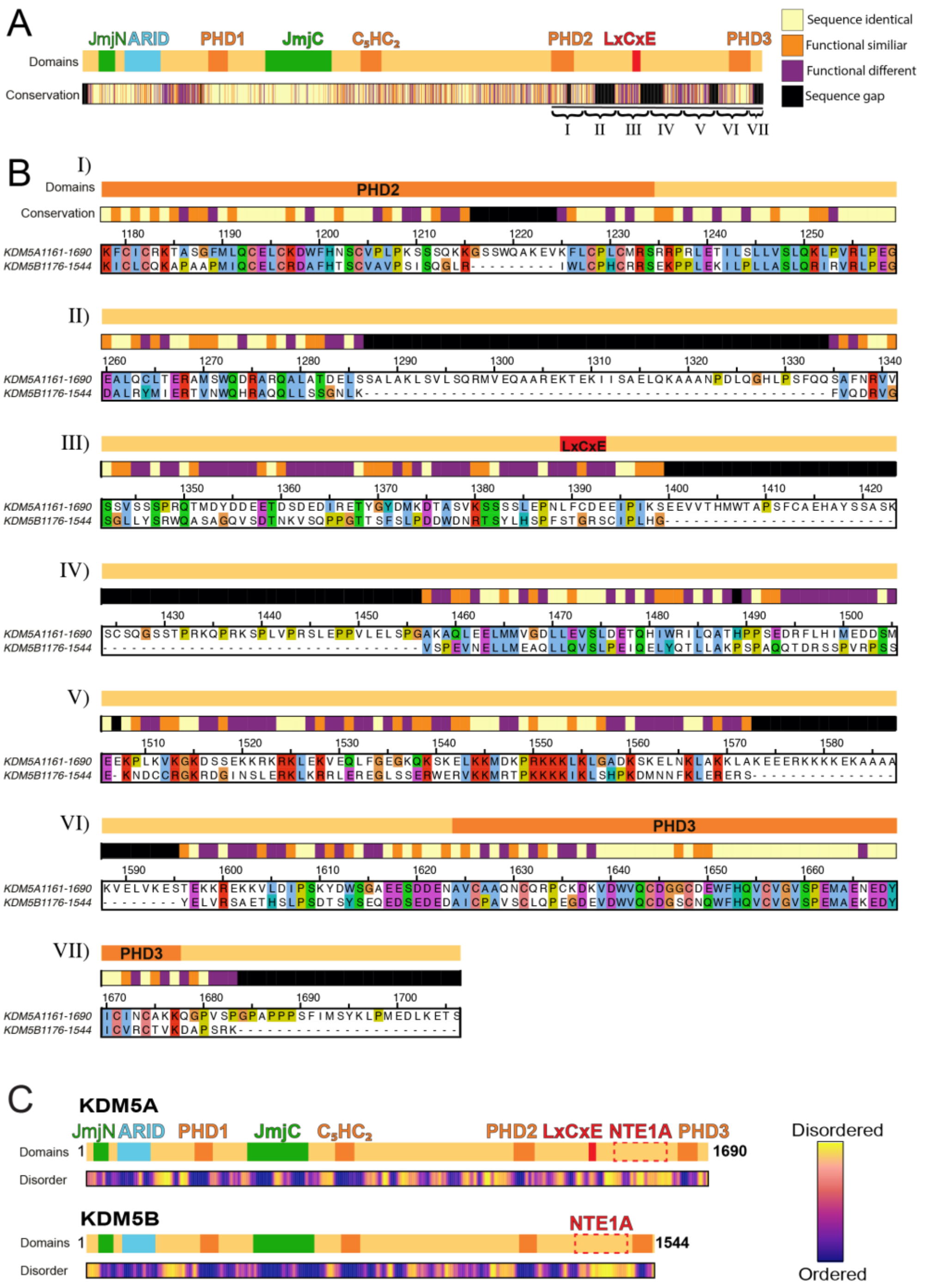


**Supplementary Figure 1. KDM5A/B Sequence alignment**

(A,B) Sequence alignment of KDM5A and KDM5B. (A) Overview of the full-length alignment, with a schematic domain map above and conservation plot below. (B), Detailed view of the sequence sections numbered in Roman letter in (A), encompassing the PHD2 domain to the C-terminus (KDM5A: aa1161-1690, KDM5B: aa1176-1544). In addition to the domain overview and the sequence plot, the aligned aa sequences of KDM5A and KDM5B are shown. A conservation score of >7 was assumed to be a functionally different amino acid, and a score of <7 was counted as a functionally similar amino acid. (C) Predicted disorder of KDM5A (top) and KDM5B (bottom) shown alongside their domain orientation. Yellow and blue colors correspond to high and low predicted disorder, respectively.

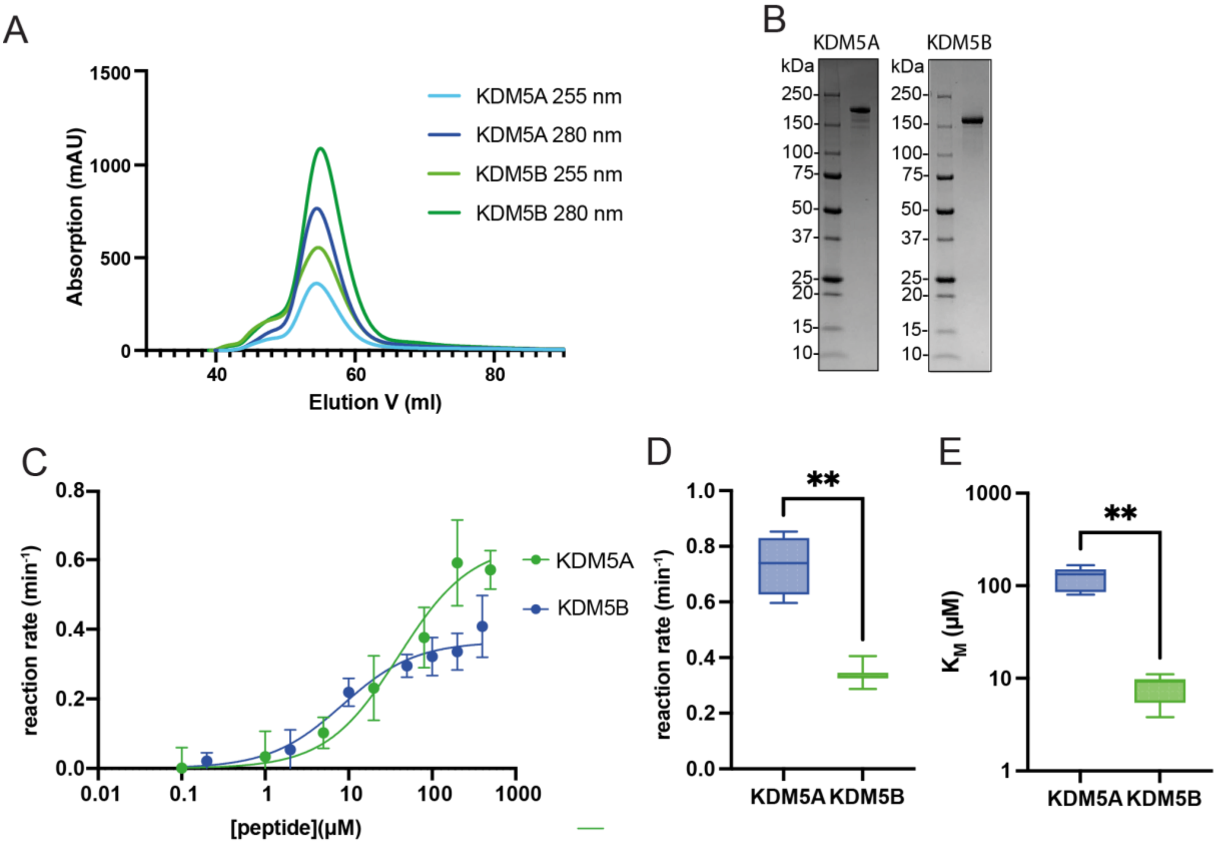


**Supplementary Figure 2. Purification of KDM5A and KDM5B**

(A) Size exclusion chromatogram of KDM5A (blue) and KDM5B (green) purification on a HiLoad 16/600 Superdex 200 pg column (Cytiva). (B) SDS-PAGE of purified KDM5A (left) and KDM5B (right) analyzed on a 5-20% gradient polyacrylamide gel stained with Coomassie Brilliant Blue. 5 µl of 2 µM KDM5 proteins were loaded per lane. (C-E) Demethylase assays of KDM5A and B using $H3_{1-18}K4me3$ peptide using the Succinate-Glo™ JmjC Demethylase/Hydroxylase Assay from Promega. (C) Michaelis–Menten plot and (D) respective calculated kinetic rate in $min^{-1}$ and (E) $K_M$ in µM of WT KDM5A (blue), WT KDM5B (green). Means of at least four experiments and the standard error of the mean are shown. Significance was determined using one-way ANOVA, with $P < 0.05$ considered significant. P-values below 0.01 are highlighted with two asterisks (**).

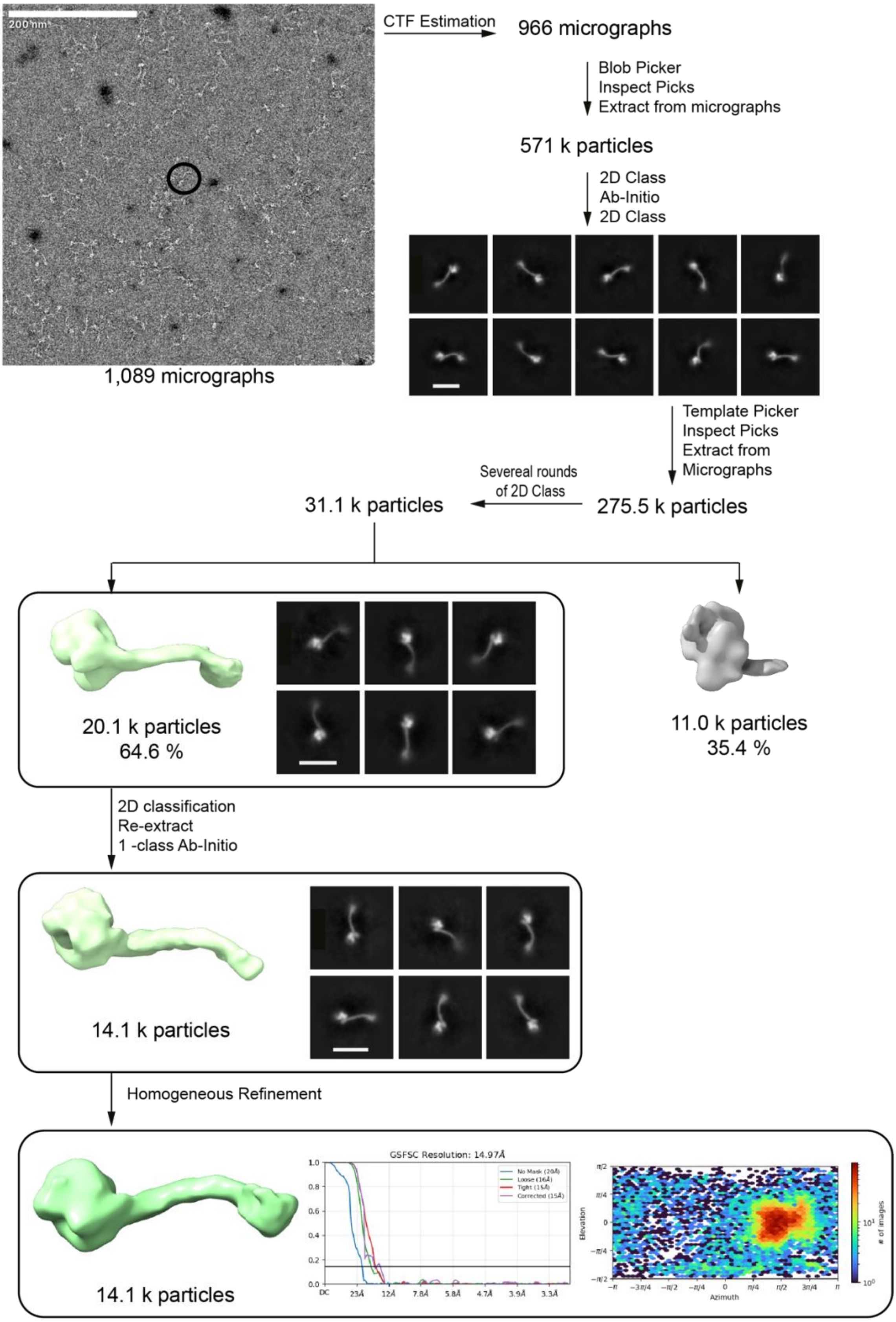


**Supplementary Figure 3. Single-particle negative-staining EM image-processing pipeline for KDM5A.** Flow chart of classifications and refinements, resulting in reconstructions of KDM5A (A) and KDM5B (B). Boxes indicate 3D reconstructions that were further used for molecular modeling. The number of particles used in each reconstruction is indicated. 2D classifications of those particle stacks yielded the shown 2D class averages. The scale bar size represents 240 Å.

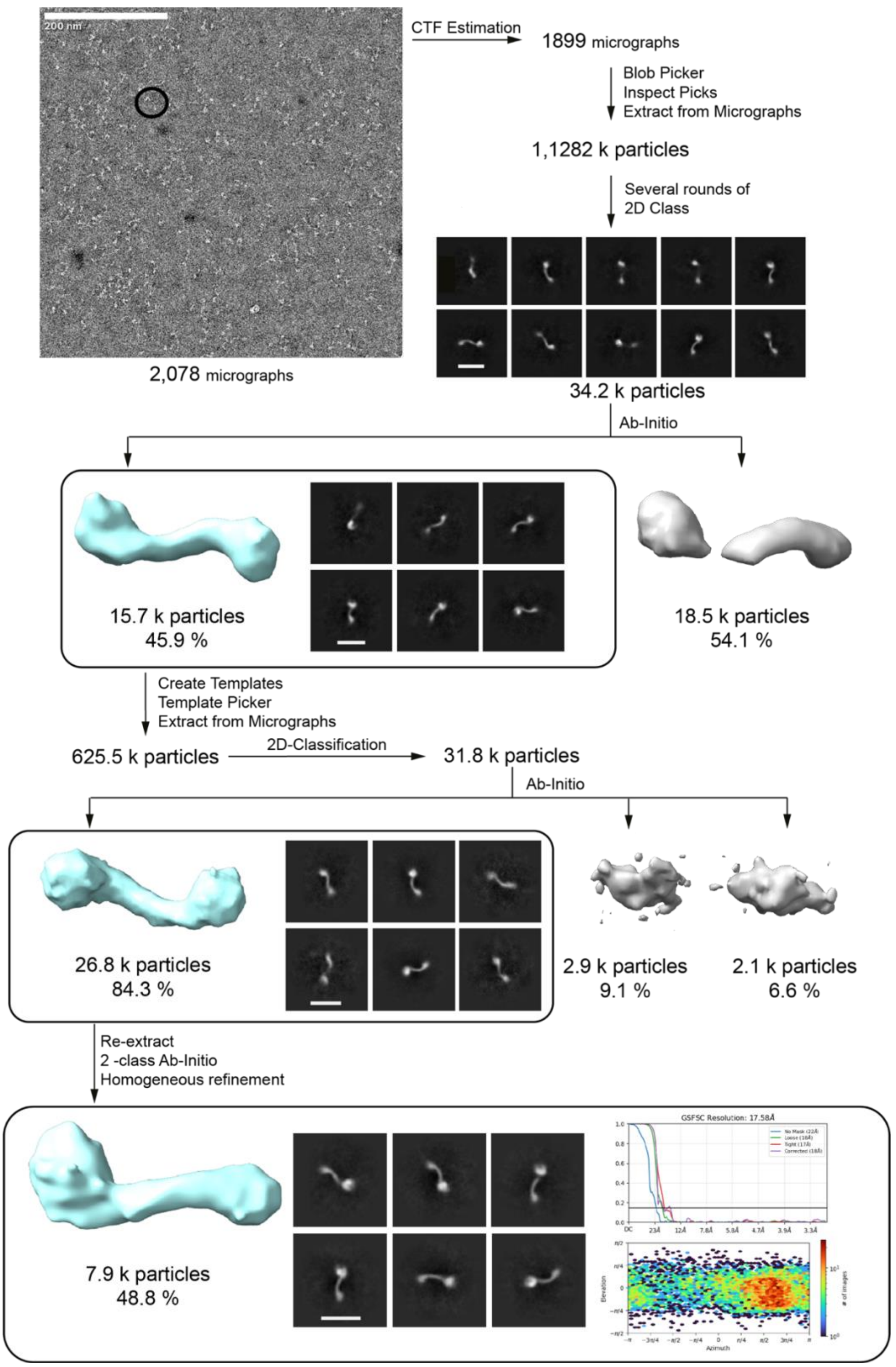


**Supplementary Figure 4. Single-particle negative-staining EM image-processing pipeline for KDM5B**

Flow chart of classifications and refinements, resulting in reconstructions of KDM5A (A) and KDM5B (B). Boxes indicate 3D reconstructions that were further used for molecular modeling. The number of particles used in each reconstruction is indicated. 2D classifications of those particle stacks yielded the shown 2D class averages. The scale bar represents 240 Å.

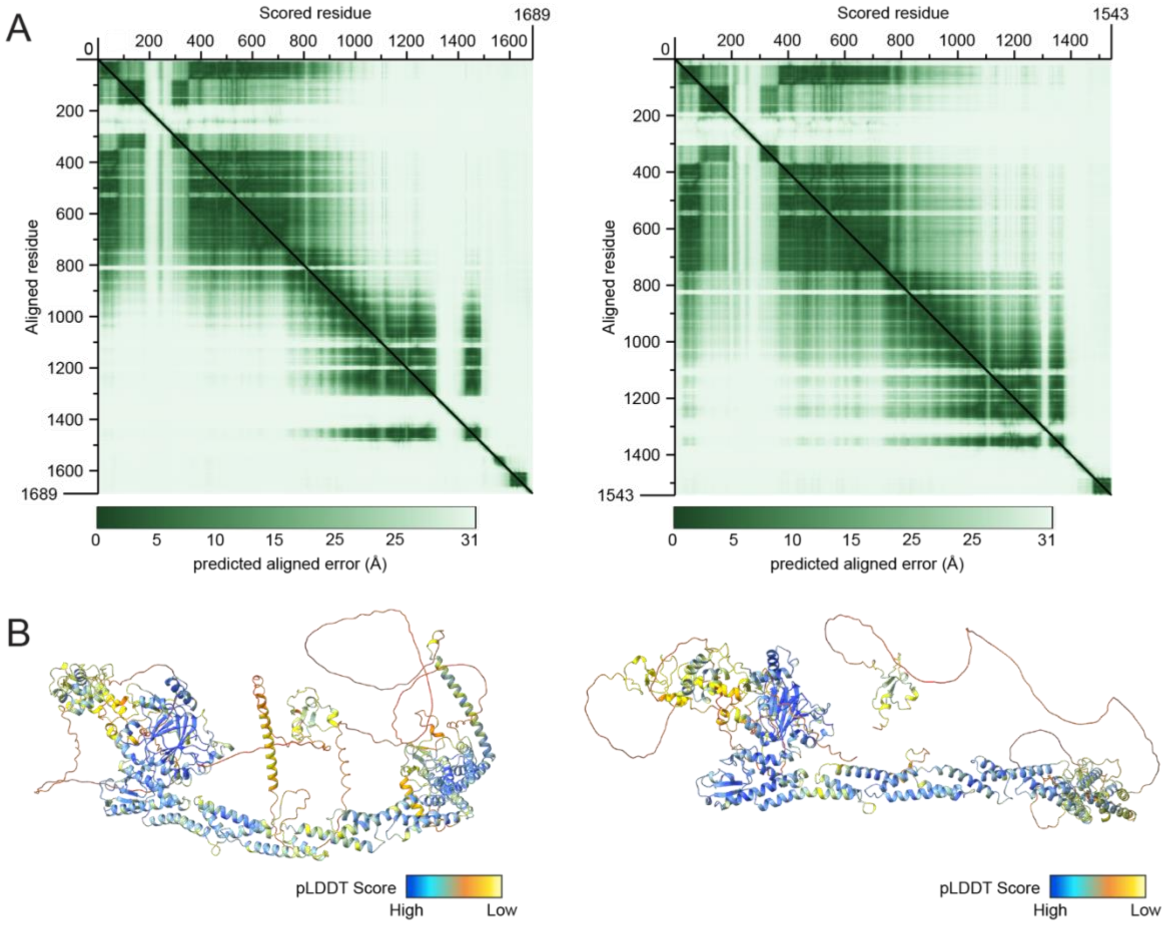


**Supplementary Figure 5. KDM5A/B Structure predictions**

(A) Predicted aligned error (pAE) score of an AlphaFold3 prediction of KDM5A (left) and KDM5B (right). The pAE scale bar is displayed below the image (B) Ribbon diagram of KDM5A (left) and KDM5B (right) AlphaFold structure prediction, colored according to pLDDT (predicted Local Distance Difference Test) confidence scores, ranging from 0 to 100, with a color gradient from orange (0 = low confidence) to blue (100 = high confidence), as highlighted in the pLDDT scale below.

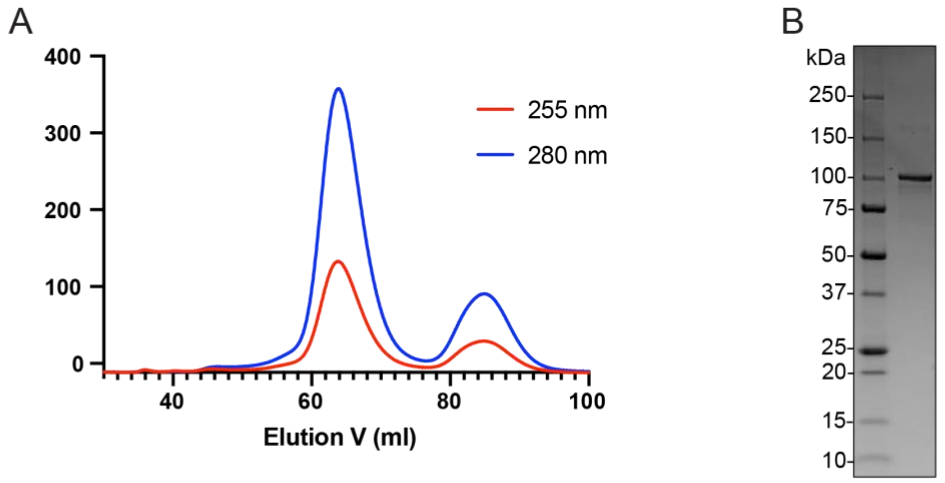


**Supplementary Figure 6. Purification of RB**

(A) Size exclusion chromatogram of WT RB purification on a HiLoad® 16/600 Superdex® 200 pg column (Cytiva) using an NGC system by BioRad. (B) SDS-PAGE of purified RB analyzed on a 5-20% gradient polyacrylamide gel stained with Coomassie Brilliant Blue. 5 µl of 2 µM RB was loaded on the gel.

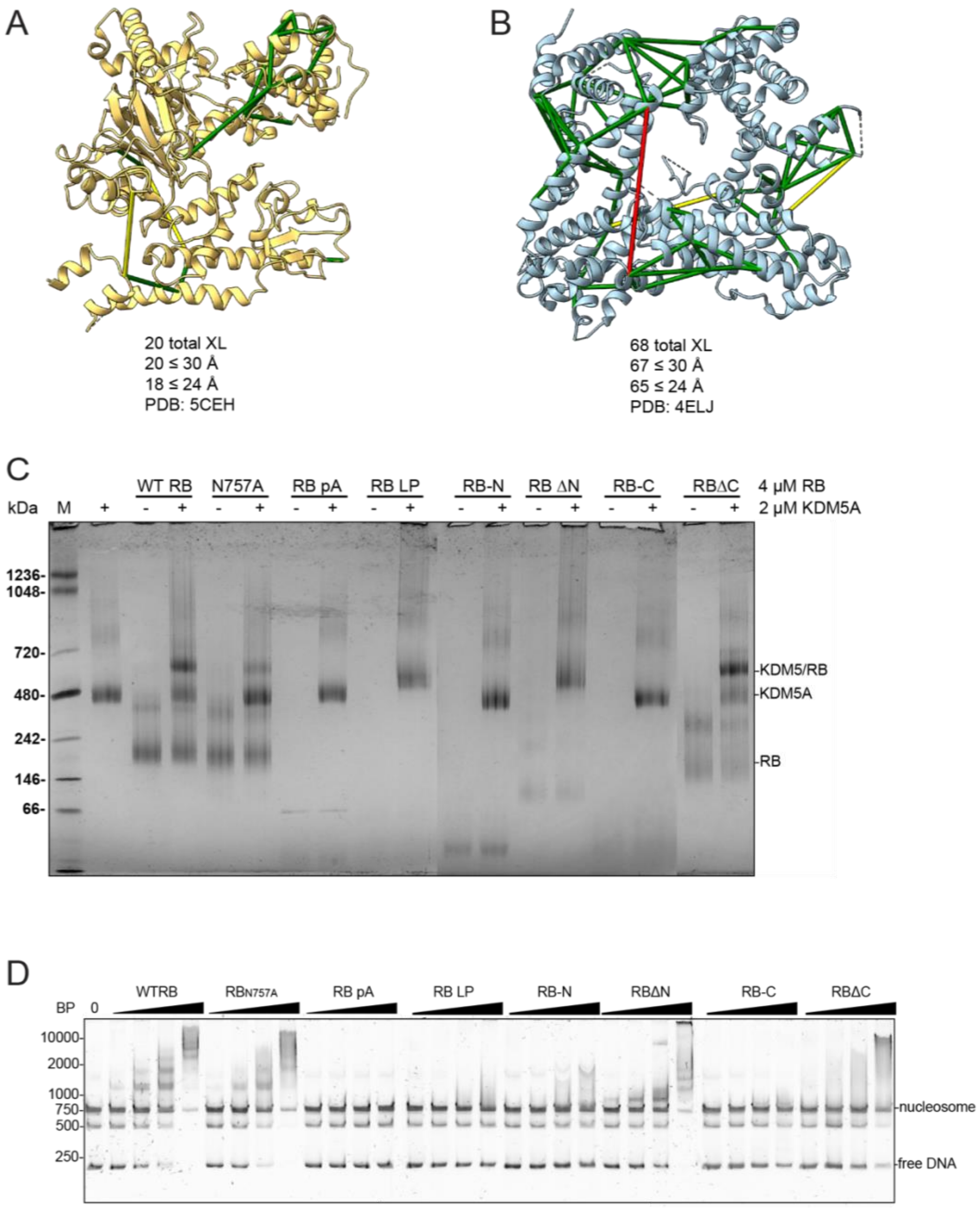


**Supplementary Figure 7. Cross-linking validation and blue native PAGE**

Validation of intramolecular cross-links indicated by lines shown in the structure of (A) truncated KDM5A (PDB:5CEH) and (B) truncated RB (PDB: 4ELJ). Cross-links shorter than 24 Å are shown in green, cross-links between a length of 24 - 30 Å are shown in yellow, and cross-links above 30 Å are shown in red. The distances between the Cα atoms of the identified cross-linked amino acids in the PDB were measured using ChimeraX. (C) Blue native PAGE of WT KDM5A incubated with either WT RB or LxCxE binding-deficient mutant of RB ($RB_{N757A}$), RB pocketA domain (RB pA), RB large pocket domain (RB LP), RB-N domain (RB-N), deletion of RB-N domain (RBΔN), RB-C region (RB-C), or deletion of RB-C region (RBΔC) (N≥3). (D) EMSA of 100 nM H3K4me3MLA nucleosomes with linker DNA incubated with 0-400 nM of either WT RB, RB N757A, RB pA, RB LP, RB-N, RBΔN, RB-C, or RBΔC (N≥3). DNA was stained with SybrGold. Sizes of fragments are indicated in base pairs (BP). The image is representative of three independent experiments.

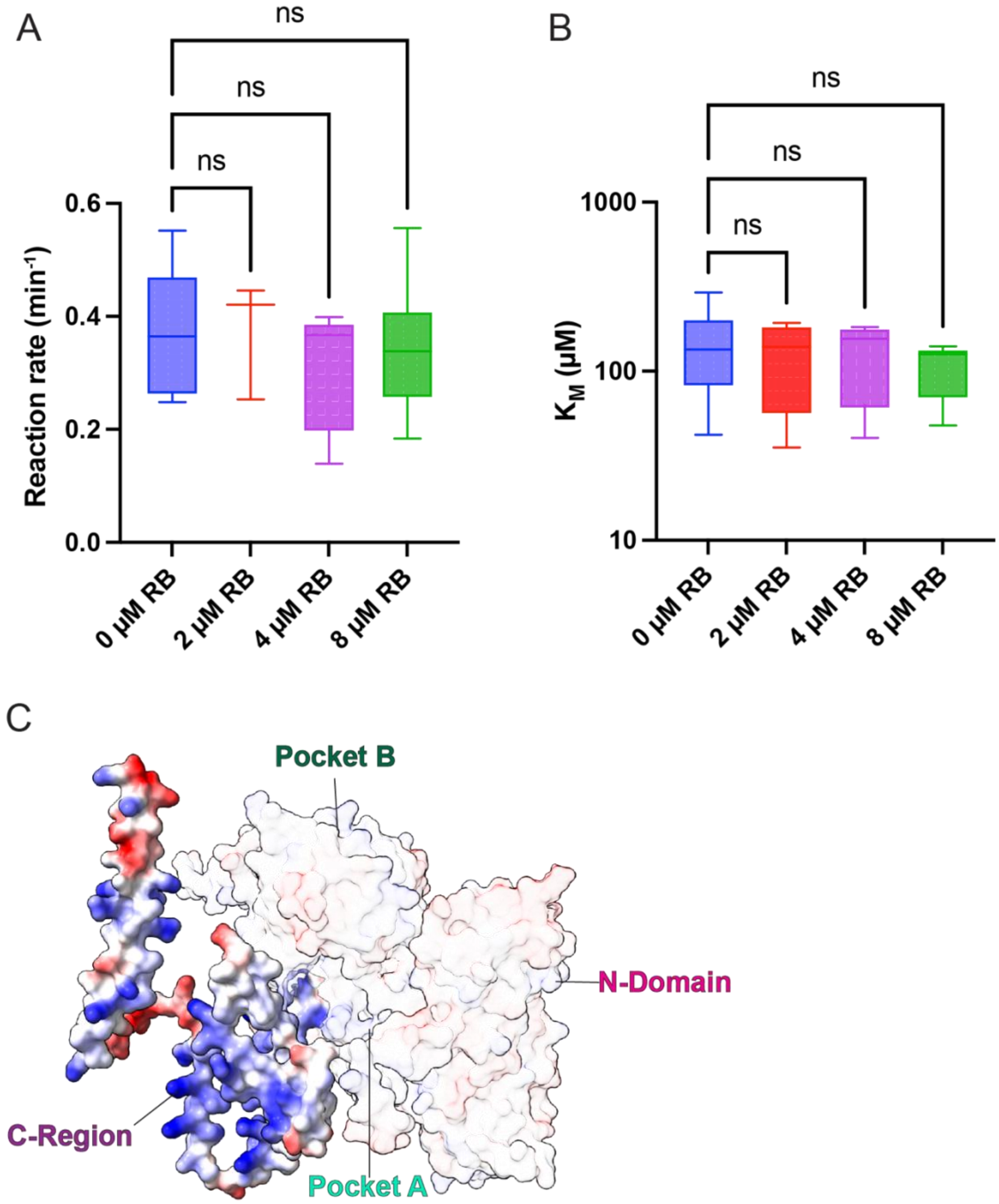


**Supplementary Figure 8. Impact of RB on peptide demethylation by KDM5A and Electrostatic surface potential of RB.**

FDH-based demethylase activity assays of KDM5A with $H3_{1\text{-}18}K4me3$ peptides as substrate under the addition of RB. Michaelis–Menten parameters with calculated kinetic rate in $min^{-1}$ (A) and $K_M$ in µM (B). Michaelis-Menten parameters were determined by direct fitting using Graphpad. Means of at least four experiments and the standard error of the mean are shown. Significance was determined using one-way ANOVA, with $P < 0.05$ considered significant. (C) Surface representation of RB colored by electrostatic potential, calculated using ChimeraX based on AlphaFold prediction available on the AlphaFold Protein Structure Database (AF-P06400-F1-v6). Blue regions indicate positively charged surfaces, red regions indicate negatively charged surfaces, and white regions represent neutral or hydrophobic areas. Domains are indicated, and the RB-C domain is highlighted.

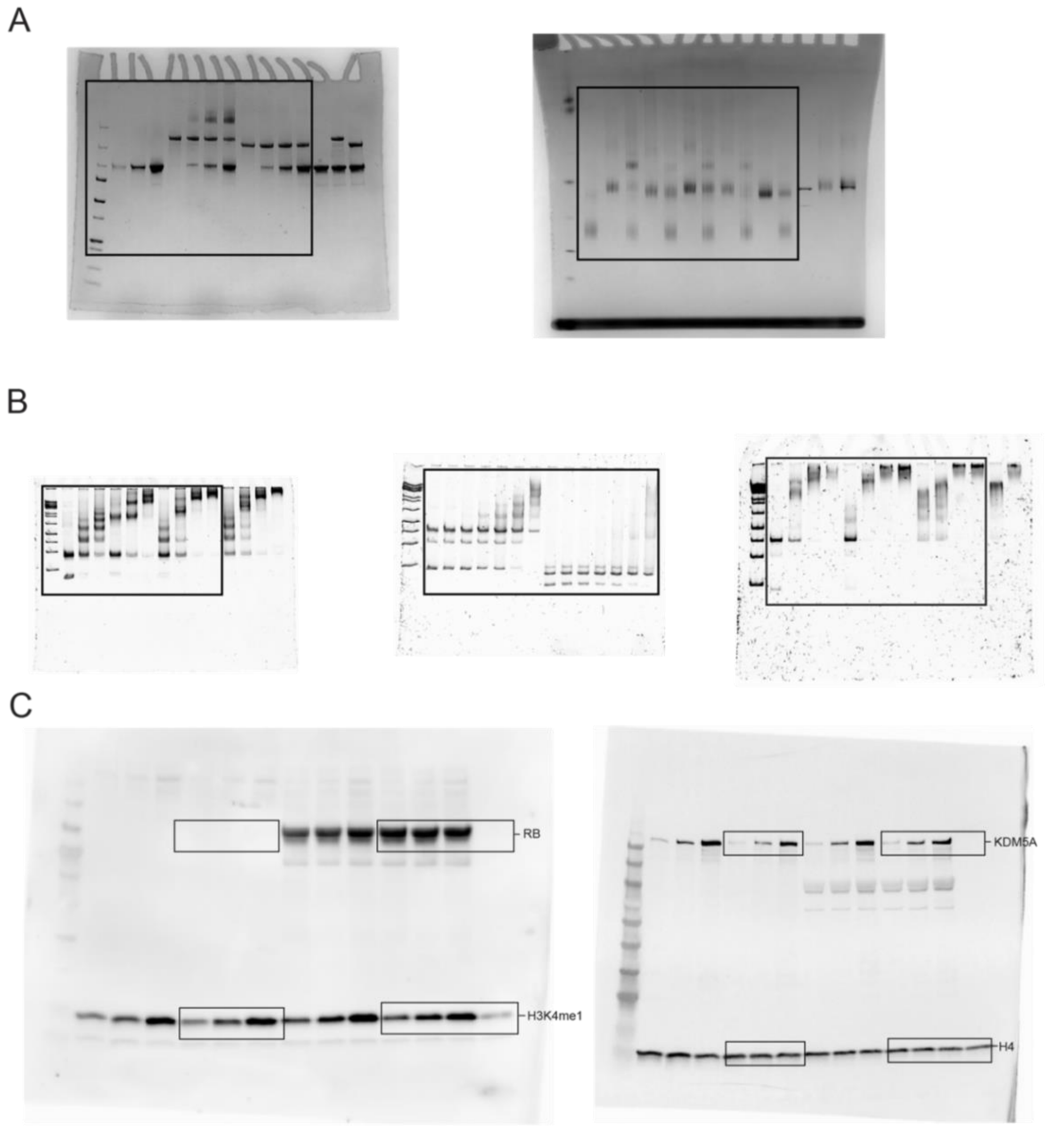


**Supplementary Figure 9. Uncropped gel images**

(A) Uncropped cross-linking gel (left) and blue native PAGE from figure 2A/B. Gels were stained using Coomasie brilliant blue (B). Uncropped EMSA gel with SybrGold staining shown in (left) figure 4A, (middle) figure 4B, and (right) figure 4C. (C) Western blot of KDM5A demethylase activity on H3K4me3MLA nucleosome, (left) RB (unspecific HRP coupled signal, due to amount of RB) and H3K4me1 imaged by HRP coupled luminescence. (right) KDM5A and H4 were imaged using fluorescence-coupled antibodies. Images represent at least three independent experiments.
