## Supplementary material for "Differences in substrate engagement and Retinoblastoma protein (RB) binding of human KDM5A and KDM5B": Sample_Legend_and_LC-MS_Settings_v01.docx.

### Overview

| **ACE project** | **Title** |
| --- | --- |
| ACE_0945 | Crosslinking of KDM5A with RB usingDSBU as a crosslinker |
| ACE_0950 | Crosslinking of KDM5A with RB usnig BS3 as a crosslinker |

### ACE_0945

#### File legend

| **ACE ID** | **Organism** | **Organ/ cell line** |  | **Treatment/ experimental setup** |
| --- | --- | --- | --- | --- |
| ACE_0945_TR01 | TRMS01_ISD | human protein, Trioplusia ni cells | Recomb. protein | Recombinant proteins, Purified by strep pulldown followed by SEC, stored in B#_B, chemically cross-linked with DSBU in B#_C quenched with 50 mM NH_4_CO_3_H. For insolution digest |
| ACE_0945_TR02 | TRMS02_Gel | human protein, Trioplusia ni cells | Recomb. protein | Recombinant proteins, Purified by strep pulldown followed by SEC, stored in B#_B, chemically cross-linked with DSBU in B#_C quenched with 50 mM NH_4_CO_3_H. SDS_Page gel cutout, for in gel digest |

#### LC_Settings

| MS device | Orbitrap Fusion Lumos |
| --- | --- |
| LC device | Thermo Vanquish Neo |
| ion source | Thermo Nanospray Flex |
| **Analytical column** | Self-packed fused silica capillary with an integrated sintered frit; CoAnn Technologies ICT36007515F-50-5 |
| column diameter | Length (L_C_) = 28 cm; ID = 75µm; OD = 360 µm; emitter 15 µm |
| stationary phase | Phenomenex Kinetex C18-XB core shell |
| particle diameter (d_p_) | 1.7 µm |
| Pore size | 120 Å |
| Column ID | AC159 |
| Column oven | Sonation column oven PRSO-V2 |
| Column oven temp. | 50°C |
| **solvents** | A: 0.2% FA, 2% ACN, 98% H_2_O  B: 0.2% FA, 80% ACN, 20 % H_2_O |
| gradient | 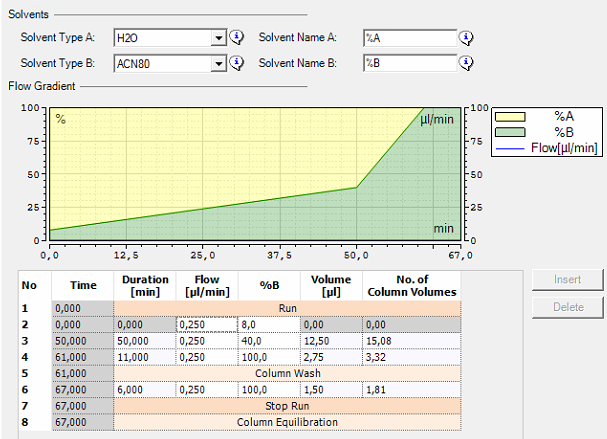 |

#### MS_Settings

##### 5.2.1. Lumos

| **Project** | **MS** | **general** | **MS1** | **MS2** | **MS2** | **MS3** | **Comments; special settings** |
| --- | --- | --- | --- | --- | --- | --- | --- |
| ACE_0945 | Lumos | Tune v4.1.4244  Xcalibur v4.7.69.37  SII: 1.7.0.468  Gradient: 67 | Analyzer: FT  Res.: 60000  SR: 380 - 1400  AGC: Standard  AGC abs.: 400000  AcT: Auto  RF: 30  SF: --  DDM: CT/3sec | Analyzer: FT  Res./ScR: 30000/-  SR: Auto  AGC: 300%  AGC abs.: 150000  AcT: 70 ms  CS: +3 to +8  IsM: Q  IsW: 1.6  Frag.: sHCD  NCE: 25, 30, 40 |  |  | classic orbitrap experiment: MS1 in Orbitrap at high resolution and data dependent MS2 also in Orbitrap high resolution. Dynamic exclusion enabled (exclude after n times=1; Exclusion duration (s)= 30; mass tolerance= ± 10ppm)  Intensity Threshold: 50000  Ion transfer Tube Temp: 270 °C  Ion Source Voltage: 2300 V |

Note: **FT**= Fourier Transform (Orbitrap); **IT**= Iontrap; **Q**= Quadrupol; **Res.**= max. Resolution at 200 m/z (Lumos) or 400 m/z (Elite) [FWHM (full width at half maximum)]; **ScR**= scan rate for measurements in the IT; **SR**= scan range [m/z]; **AGC**= automatic gain control, max number of acquired ions per measurement; **AcT**= max. Ion acquisition time [ms]; **CS**= charge states used for fragmentation; **IsM**= Isolation mode (Q or IT), MS2 isolation and further is only done in IT; **IsW**= Isolation window [m/z], value followed by scan mode the isolation is based on (MS1, MS2 …) **Frag.**= Fragmentation method; **HCD**= Higher-energy collisional dissociation; **CID**= Collision-induced dissociation; **ETD**= Electron-transfer dissociation**; EThcD=** Electron-Transfer/Higher-Energy Collision Dissociation; **sHCD**= stepped HCD**; NCE**= normalized collision energy; **cycles**: number of MSn recorded or max cycle time; RF= RF Lens [%]; **SF**= Source Fragmentation [V]; **DDM**: Data dependent Mode (cycle time in seconds, CT/[s] or number of scans, NS); **NS**= Number of data dependent scans

#### Search Settings

| Program & version | MetaMorpheus v1.0.6. |
| --- | --- |
| Search engine | MetaMorpheusXL |
| settings | 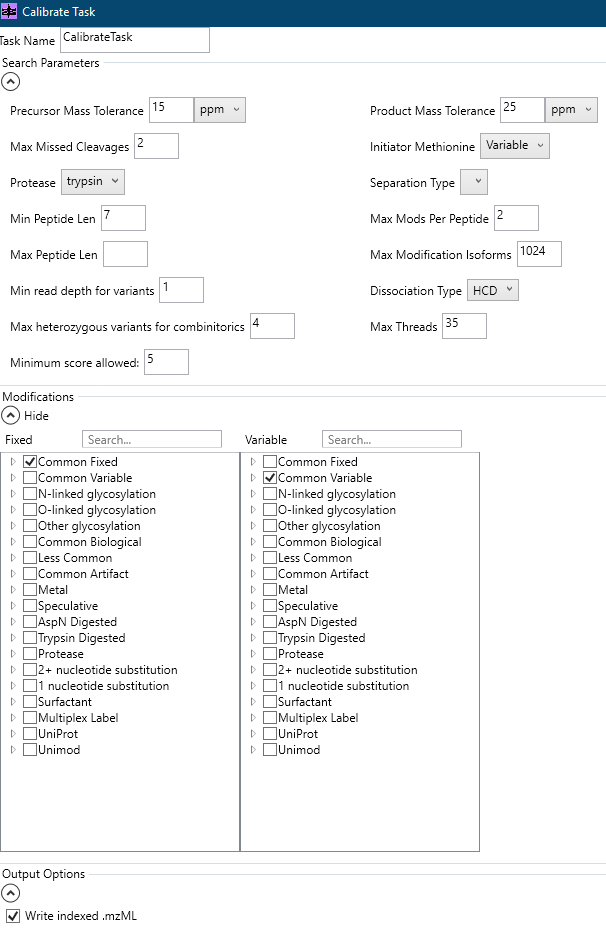 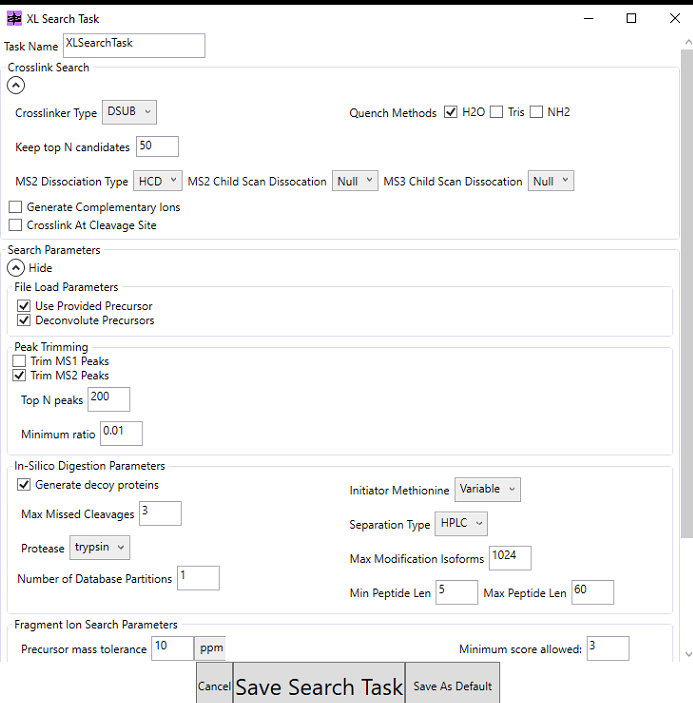 |
| Static modification | Carbamidomethyl (C) |
| Digestion mode | Trypsin/P (specific), 3 missed cleavages |
| CL and specificity | DSBU; Site A: **K**; Site B: **K or KSTY** |
| Dynamic modification | Oxidation (M) |
| Program & version | 1. ACE_0945_0950_SOI_plus_con_v01.fasta |
| Annotation |  |

Note: Database has contaminants appended.

### ACE_0950

#### File legend

| **ACE ID** | **Organism** | **Organ/ cell line** |  | **Treatment/ experimental setup** |
| --- | --- | --- | --- | --- |
| ACE_0950_TR03 | TRMS01_ISD | human protein, Trioplusia ni cells | Recomb. protein | Recombinant proteins, Purified by strep pulldown followed by SEC, stored in B#_B, chemically cross-linked with DSBU in B#_C quenched with 50 mM NH_4_CO_3_H. For insolution digest |
| ACE_0950_TR04 | TRMS02_Gel | human protein, Trioplusia ni cells | Recomb. protein | Recombinant proteins, Purified by strep pulldown followed by SEC, stored in B#_B, chemically cross-linked with DSBU in B#_C quenched with 50 mM NH_4_CO_3_H. SDS_Page gel cutout, for in gel digest |

#### LC_Settings

| MS device | Orbitrap Fusion Lumos |
| --- | --- |
| LC device | Thermo Vanquish Neo |
| ion source | Thermo Nanospray Flex |
| **Analytical column** | Self-packed fused silica capillary with an integrated sintered frit; CoAnn Technologies ICT36007515F-50-5 |
| column diameter | Length (L_C_) = 28 cm; ID = 75µm; OD = 360 µm; emitter 15 µm |
| stationary phase | Phenomenex Kinetex C18-XB core shell |
| particle diameter (d_p_) | 1.7 µm |
| Pore size | 120 Å |
| Column ID | AC159 |
| Column oven | Sonation column oven PRSO-V2 |
| Column oven temp. | 50°C |
| **solvents** | A: 0.2% FA, 2% ACN, 98% H_2_O  B: 0.2% FA, 80% ACN, 20 % H_2_O |
| gradient | 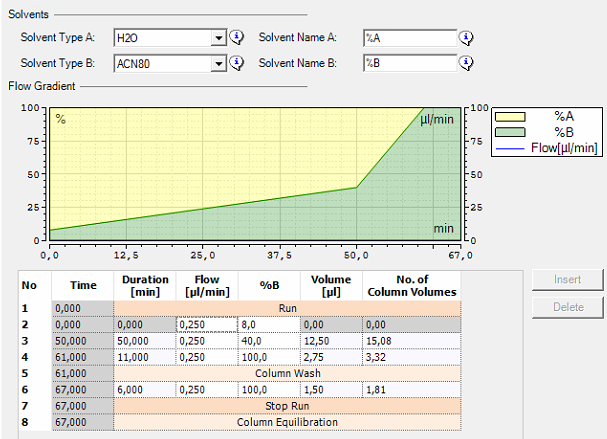 |

#### MS_Settings

| **Project** | **MS** | **general** | **MS1** | **MS2** | **MS2** | **MS3** | **Comments; special settings** |
| --- | --- | --- | --- | --- | --- | --- | --- |
| ACE_0950 | Lumos | Tune v4.1.4244  Xcalibur v4.7.69.37  SII: 1.7.0.468  Gradient: 67 | Analyzer: FT  Res.: 60000  SR: 380 - 1400  AGC: Standard  AGC abs.: 400000  AcT: Auto  RF: 30  SF: --  DDM: CT/3sec | Analyzer: FT  Res./ScR: 30000/-  SR: Auto  AGC: 300%  AGC abs.: 150000  AcT: 70 ms  CS: +3 to +8  IsM: Q  IsW: 1.6  Frag.: sHCD  NCE: 25, 30, 40 |  |  | classic orbitrap experiment: MS1 in Orbitrap at high resolution and data dependent MS2 also in Orbitrap high resolution. Dynamic exclusion enabled (exclude after n times=1; Exclusion duration (s)= 30; mass tolerance= ± 10ppm)  Intensity Threshold: 50000  Ion transfer Tube Temp: 270 °C  Ion Source Voltage: 2300 V |

Note: **FT**= Fourier Transform (Orbitrap); **IT**= Iontrap; **Q**= Quadrupol; **Res.**= max. Resolution at 200 m/z (Lumos) or 400 m/z (Elite) [FWHM (full width at half maximum)]; **ScR**= scan rate for measurements in the IT; **SR**= scan range [m/z]; **AGC**= automatic gain control, max number of acquired ions per measurement; **AcT**= max. Ion acquisition time [ms]; **CS**= charge states used for fragmentation; **IsM**= Isolation mode (Q or IT), MS2 isolation and further is only done in IT; **IsW**= Isolation window [m/z], value followed by scan mode the isolation is based on (MS1, MS2 …) **Frag.**= Fragmentation method; **HCD**= Higher-energy collisional dissociation; **CID**= Collision-induced dissociation; **ETD**= Electron-transfer dissociation**; EThcD=** Electron-Transfer/Higher-Energy Collision Dissociation; **sHCD**= stepped HCD**; NCE**= normalized collision energy; **cycles**: number of MSn recorded or max cycle time; RF= RF Lens [%]; **SF**= Source Fragmentation [V]; **DDM**: Data dependent Mode (cycle time in seconds, CT/[s] or number of scans, NS); **NS**= Number of data dependent scans

#### Search Settings

| Program & version | MetaMorpheus v1.0.6. |
| --- | --- |
| Search engine | MetaMorpheusXL |
| settings | 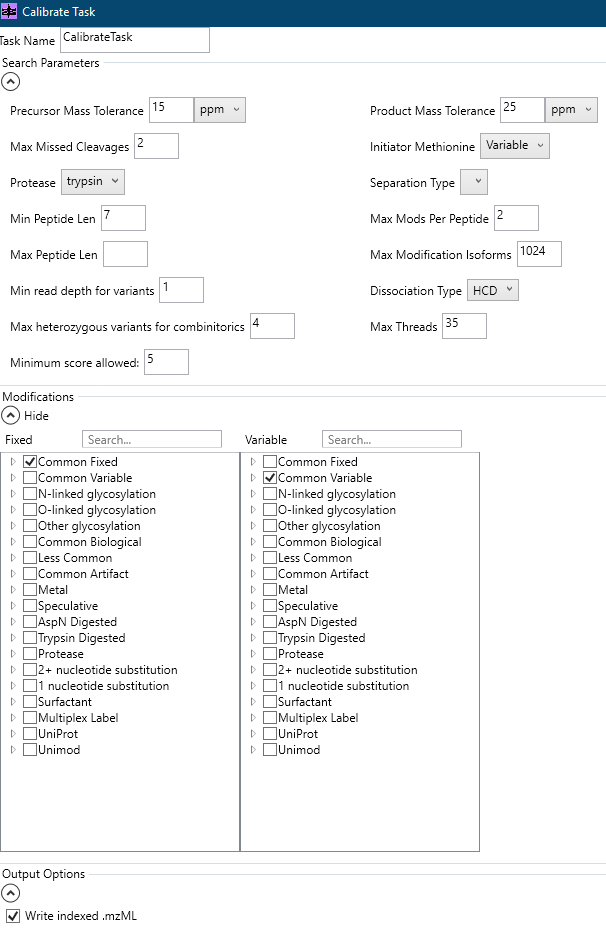 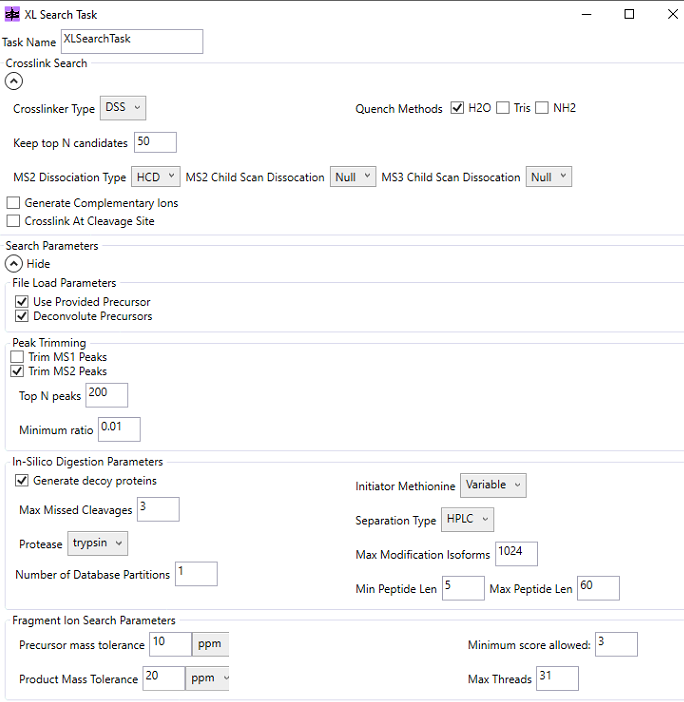 |
| Static modification | Carbamidomethyl (C) |
| Digestion mode | Trypsin/P (specific), 3 missed cleavages |
| CL and specificity | BS^3^; Site A: **K**; Site B: **K** |
| Dynamic modification | Oxidation (M) |
| Program & version | 1. ACE_0945_0950_SOI_plus_con_v01.fasta |
| Annotation |  |

Note: Database has contaminants appended. Note DSS and BS3 have same setting.
